# Gamma-Mobile-Trio systems define a new class of mobile elements rich in bacterial defensive and offensive tools

**DOI:** 10.1101/2023.03.28.534373

**Authors:** Tridib Mahata, Katarzyna Kanarek, Moran G. Goren, Marimuthu Ragavan Rameshkumar, Eran Bosis, Udi Qimron, Dor Salomon

**Affiliations:** Department of Clinical Microbiology and Immunology, School of Medicine, Faculty of Medical and Health Sciences, Tel Aviv University, Tel Aviv, Israel; Department of Biotechnology Engineering, Braude College of Engineering, Karmiel, Israel

**Keywords:** T6SS, anti-phage, toxin, effector, antibacterial, nuclease, DNA mobility, integrase, GMT, Vibrio

## Abstract

Conflicts between bacteria and their rivals led to an evolutionary arms race and the development of bacterial immune systems. Although diverse immunity mechanisms were recently identified, many remain unknown, and their dissemination within bacteria is poorly understood. Here, we describe a widespread genetic element, defined by the presence of the Gamma-Mobile-Trio (GMT) proteins, that serves as a bacterial survival kit. We show that GMT-containing genomic islands are active mobile elements with cargo comprising various anti-phage defense systems, in addition to antibacterial type VI secretion system (T6SS) effectors and antibiotic resistance genes. We identify four new anti-phage defense systems encoded within GMT islands. A thorough investigation of one system reveals that it is triggered by a phage capsid protein to induce cell dormancy. Our findings underscore the need to broaden the concept of ‘defense islands’ to include also antibacterial offensive tools, such as T6SS effectors, as they share the same mobile elements as defensive tools for dissemination.

## Introduction

Competition and predation contribute to bacterial evolution. For example, the type VI secretion system (T6SS), an offensive, missile-like dueling apparatus that delivers antibacterial toxins (i.e., effectors) directly into rival bacteria^1–5^, was shown to shift the balance of bacterial populations and lead to the emergence of dominant strains^6–11^. Similarly, bacteriophages (phages) drive bacterial evolution and population dynamics^12–14^. Their ability to prey on bacteria has led to an arms race in which bacteria evolve or acquire anti-phage defense systems while phages counteract these actions with anti-defense mechanisms^15^. Interestingly, anti-phage defense systems often cluster in so-called ‘defense islands’^16,17^, a phenomenon that has been used to identify dozens of anti-phage defense systems in recent years^18–20^. Probably, many defense systems are yet to be revealed. Identifying additional defense systems or antibacterial toxins and deciphering the mechanisms governing their spread within bacterial populations is important to understanding bacterial evolution.

Mobile genetic elements (MGEs), such as plasmids, phages, and transposons, mediate horizontal gene transfer (HGT) and play a significant role in bacterial evolution by enhancing bacterial fitness^21–27^. Anti-phage defense systems, secreted antibacterial toxins, and antibiotic-resistance genes have been identified within MGEs and were predicted to be horizontally shared^13,20,28–35^. Nevertheless, many MGEs are poorly understood and require further investigation to reveal their distribution mechanisms and cargo.

We previously reported that *Vibrio parahaemolyticus* island-6 (VPaI-6), a genomic island encompassing *vpa1270*-*vpa1254* on chromosome 2 of the human pathogen *V. parahaemolyticus* RIMD 2210633 (hereafter referred to as RIMD)^36^, encodes an antibacterial DNase T6SS effector, VPA1263 (BAC62606.1), and its cognate immunity protein, Vti2 (WP_005477334.1)^37,38^. Here, we describe a protein trio found in VPaI-6 and other genomic islands, which defines a widespread mobile genetic element with diverse cargo rich in antibacterial T6SS effectors and anti-phage defense systems. Examining genes and operons of unknown function within these islands revealed four new anti-phage defense systems. Therefore, the described MGE is akin to a bacterial armory containing diverse offensive and defensive tools against potential rivals, which can be horizontally shared.

## Results

### GMT proteins define a new class of genomic islands

VPaI-6 is found in *V. parahaemolyticus* RIMD and a subset of other *V. parahaemolyticus* strains^36^ (**Fig. 1a**). Analysis of VPaI-6 revealed that the first three genes, encoding VPA1270, VPA1269, and VPA1268 (WP_005477115.1, WP_005477239.1, and WP_005477284.1, respectively), are annotated in the NCBI protein family models database as a Gamma-Mobile-Trio (GMT) system. We found homologous co-occurring GMT proteins encoded in thousands of Gram-negative and Gram-positive bacterial genomes (**Fig. 1b** and **Dataset S1**). Since no information was available on the function of these proteins^36^, we set out to investigate the GMT system.

**Fig. 1.**
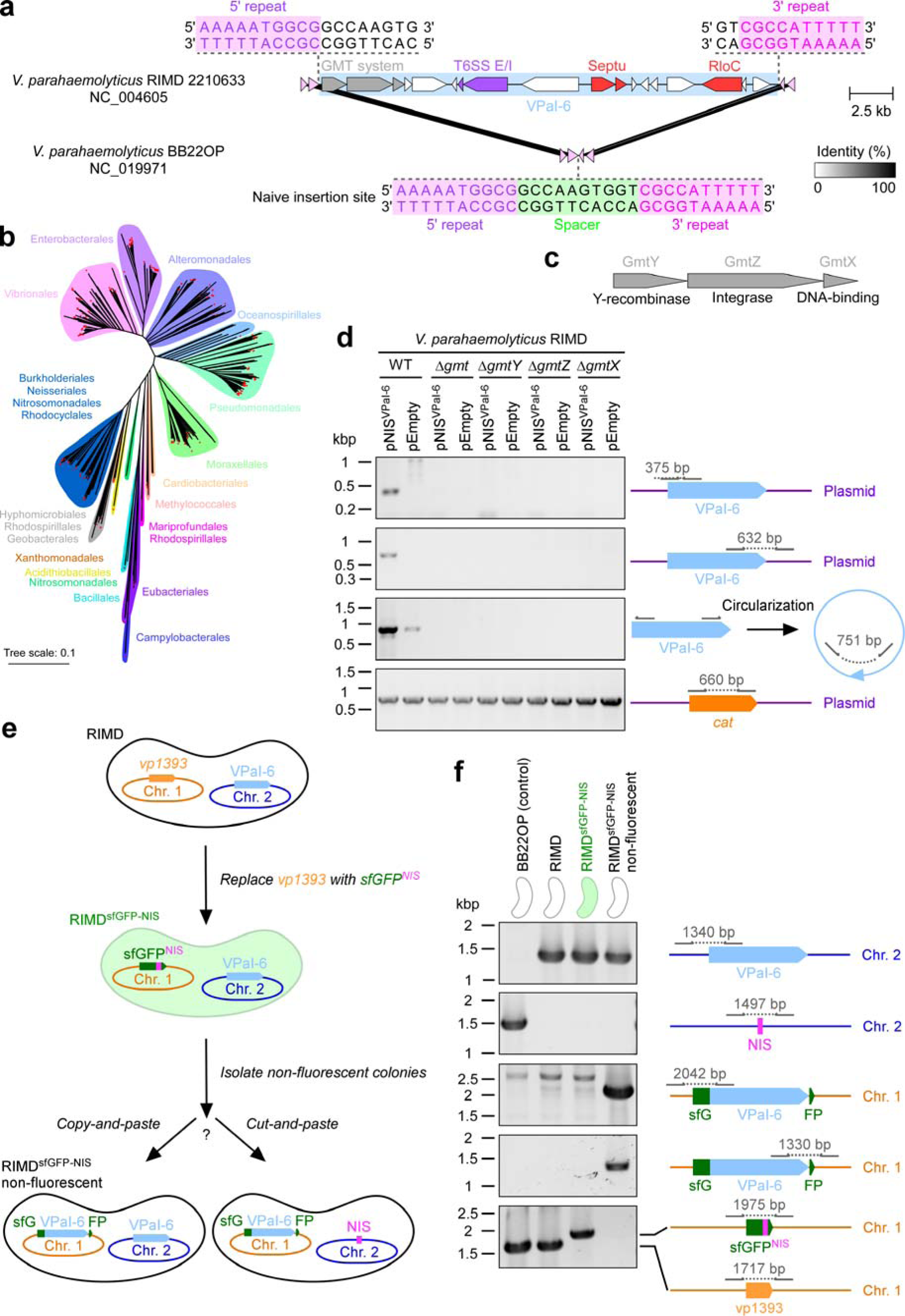
GMT proteins define a class of widespread, mobile genomic islands. **(a)** Schematic representation of VPaI-6 (cyan rectangle) and flanking regions. A predicted naïve insertion site (NIS) identified in *V. parahaemolyticus* BB22OP is shown below; gray rectangles denote protein sequence identity percentage. Inverted repeat sequences identified in the naïve insertion site and flanking VPaI-6 are denoted. RefSeq accession numbers are provided. **(b)** Phylogenetic distribution of bacteria containing GMT systems, based on the DNA sequence of *rpoB*. Bacterial orders are denoted. The evolutionary history was inferred using the neighbor-joining method. The evolutionary distances represent the number of nucleotide substitutions per site. Red stars denote bacteria in which the borders of a GMT island were determined. **(c)** Predicted activities of GMT system proteins. **(d)** Agarose gel electrophoresis analysis of the indicated amplicons. The total DNA isolated from wild-type (WT) *V. parahaemolyticus* RIMD cells or its derivative strains in which the entire GMT system was deleted (Δ*gmt*) or its individual components (i.e., Δ*gmtX*, Δ*gmtZ*, and Δ*gmtX*), conjugated with an empty plasmid (pEmpty) or a plasmid containing a predicted naïve insertion site for VPaI-6 (pNIS^VPaI-6^), was used as a template. The *cat* gene found in the backbone of both plasmids was amplified as a control for plasmid presence. **(e)** An illustration of the assay devised to distinguish between a copy-and-paste and a cut-and-paste transfer mechanism of VPaI-6. Chr. 1, chromosome 1; Chr. 2, chromosome 2; sfGFP-NIS, an sfGFP-encoding gene containing the 30 bp-long VPaI-6 NIS sequence as a linker between the 10^th^ and 11^th^ beta strands of sfGFP. **(f)** Agarose gel electrophoresis analysis of the indicated amplicons. The total DNA isolated from WT RIMD, a derivative in which *vp1393* was replaced by sfGFP-NIS (RIMD^sfGFP-NIS^), or an isolated RIMD^sfGFP-NIS^ colony that lost its fluorescence (as described in (b)), was used as a template. *V. parahaemolyticus* BB22OP was used as a control for a chromosomal VPaI-6 NIS. In (d) and (f), arrows denote the positions of primers used for each amplicon; the expected amplicon size is denoted in gray.

According to the NCBI Conserved Domain Database (CDD)^39^, the trio’s first gene, named *vpa1270* or *gmtY*, encodes a site-specific tyrosine recombinase with a domain belonging to the DNA_BRE_C superfamily. The second gene, *vpa1269*, encodes an integrase with a domain belonging to the Phage_Integr_2 superfamily, which is predicted to mediate unidirectional site-specific recombination (hereafter referred to as *gmtZ*). The third gene, named *vpa1268* or *gmtX*, encodes a protein of unknown function; we found that it is similar to the DNA binding domain of the partition protein StbA of plasmid R388 (according to HHpred^40^, ∼93% probability of similarity between amino acids 3-75 of GmtX and amino acids 1-68 of StbA [PDB: 7PC1_A]). These predicted activities suggest that the GMT system plays a role in DNA excision and integration (**Fig. 1c**).

Systematic analysis of publicly available genomes revealed that GMT systems are found in predicted genomic islands. Comparisons between GMT-containing regions identified in completely assembled genomes and the genomes of closely related bacteria suggested that GMT-encoding genes define the 5’ border of these islands. By identifying sequences that flank the GMT region but are found adjacent to each other in genomes of closely related bacteria, we further revealed the putative 3’ border for 83% (366 out of 442) of these GMT islands and their predicted naïve insertion sites (NIS; **Fig. 1a,b** and **Dataset S2**). A genome may contain multiple and diverse GMT islands, which can reside on the chromosome or a plasmid (**Dataset S1**).

Our analysis revealed diverse putative insertion sites of GMT islands in bacterial genomes, either intergenic or intragenic (**Extended Data Fig. 1** and **Dataset S2**). Notably, a phylogenetic tree of GmtY encoded within the GMT islands for which we identified a putative insertion site revealed that closely related proteins are found in diverse bacterial orders yet share a similar insertion site (**Extended Data Fig. 1**). This observation suggests that GMT islands are horizontally shared.

Interestingly, when a GMT island appears to have been inserted intragenically, we often find a homolog of the disrupted gene encoded within the island, possibly to compensate for its loss (**Extended Data Fig. 2**), as previously reported for other MGEs^13,41^. In most cases (238 of 366), we identified an inverted repeat, at least 5 nucleotides long, as part of the predicted GMT island NIS (**Dataset S2**); the same repeat is often found flanking the GMT island (**Fig. 1a**). These observations led us to hypothesize that GMT islands are mobile and that repeat sequences define specific insertion sites for each island.

### GMT islands are active mobile elements

We reasoned that if VPaI-6 is a functional MGE, it should transfer into a cognate NIS. To investigate this possibility, we introduced a low copy-number plasmid harboring a predicted 30 bp-long VPaI-6 NIS from *V. parahaemolyticus* BB22OP (pNIS^VPaI-6^; **Fig. 1a**) into wild-type RIMD cells. Using primer sets designed to amplify fusions between the ends of VPaI-6 and the plasmid sequences flanking the NIS, we found that the RIMD population indeed contained plasmids into which VPaI-6 was inserted (**Fig. 1d**). We confirmed the insertion into the plasmid-borne NIS with Sanger sequencing of the amplified products (**File S1**). Furthermore, using primers facing outward from each end of VPaI-6, we revealed the existence of a circular form of the GMT island lacking the flanking inverted repeats (**Fig. 1d** and **File S1**). The amplification products were missing when we used RIMD derivatives in which we either deleted the genes encoding the GMT system (Δ*gmt*), deleted individual GMT system components (Δ*gmtY*, Δ*gmtZ*, and Δ*gmtX*) (**Fig. 1d**), or modified the spacer sequence between the inverted repeats of the predicted NIS on the plasmid (**Extended Data Fig. 3**). Taken together, our results demonstrate that VPaI-6 is a mobile GMT island, which inserts specifically via an intermediate circular form into the repeat-containing site we identified in the above analyses.

To determine whether other GMT islands are also mobile, we investigated the ability of a GMT island found in the chromosome of *V. parahaemolyticus* 04.2548^42^, which was available in our laboratory stocks (**Extended Data Fig. 4a**), to transfer into a cognate, plasmid-borne NIS. Our results confirmed that the 04.2548 GMT island transferred into its plasmid-borne cognate NIS (pNIS^04.2548^) but not into a plasmid containing the VPaI-6 NIS (pNIS^VPaI-6^; **Extended Data Fig. 4b**). As observed with VPaI-6, we identified a circular form of the 04.2548 GMT island lacking the inverted repeats of the insertion site (**File S2**). These results indicate that GMT islands are mobile, and each GMT system identifies a specific repeat-containing sequence for insertion.

### GMT islands employ a replicative mechanism of transfer

Next, we investigated whether VPaI-6 employs a conservative or replicative mechanism of transfer (i.e., cut-and-paste or copy-and-paste, respectively) by introducing a single VPaI-6 NIS into chromosome 1 of RIMD and following the fate of VPaI-6 located on chromosome 2. To monitor the insertion of VPaI-6 into the NIS, we engineered a system in which the VPaI-6 NIS was introduced in-frame as a linker between the 10^th^ and 11^th^ β-strands of a superfolder GFP (sfGFP)^43^ to produce sfGFP^NIS^. The sfGFP^NIS^ gene was then used to replace *hcp1* (*vp1393*) on chromosome 1, resulting in GFP-fluorescent cells containing a single chromosomal copy of the VPaI-6 NIS within the sfGFP gene (RIMD^sfGFP-NIS^). To identify events of VPaI-6 insertion into the NIS, we plated the fluorescent cells and monitored the appearance of non-fluorescent colonies, suggestive of VPaI-6 transfer into the NIS, thereby obstructing the expression of a functional sfGFP protein (**Fig. 1e**). After isolating non-fluorescent cells, we confirmed that the sfGFP^NIS^ was indeed interrupted by the insertion of VPaI-6 using PCR amplification of the junctions between VPaI-6 ends and the sfGFP^NIS^ flanking sequences (**Fig. 1f**). Importantly, the original copy of VPaI-6 remained on chromosome 2. We did not identify a loss of VPaI-6 from either the original VPaI-6 location on chromosome 2 or the new location on chromosome 1 in these isolated, non-fluorescent cells. These results indicate that VPaI-6 employs a replicative transfer mechanism.

### Plasmids can mediate the horizontal transfer of GMT islands

We observed many GMT systems encoded on plasmids (**Dataset S1**), suggesting a possible plasmid-mediated horizontal transfer mechanism for these MGEs. Therefore, we sought to demonstrate that a GMT island can transfer between bacteria via a conjugatable plasmid. To this end, we constructed a RIMD derivative (RIMD^VPaI-6_Gent^) in which we replaced the VPaI-6 region encompassing *vpa1254*-*vpa1262* with a gentamicin resistance cassette (VPaI-6^Gent^; **Fig. 2a**) that enables selection of the mobilized island. A conjugatable plasmid containing the VPaI-6 NIS (pNIS^VPaI-6^) was introduced into RIMD^VPaI-6_Gent^, and the derivative GMT island was mobilized into the plasmid-borne NIS, as determined by a PCR performed on the pooled bacterial population (**Fig. 2b,c**). We then mixed this pooled population with a derivative strain of *V. parahaemolyticus* BB22OP containing a chromosomal tetracycline resistance cassette (BB22OP^Tet^), in the presence of a conjugation helper strain. BB22OP^Tet^ conjugates containing pNIS^VPaI-6^ harboring VPaI-6^Gent^ were selected on agar plates supplemented with the appropriate antibiotics, and PCR analyses revealed that these resulting colonies comprised a mixture of cells in which VPaI-6^Gent^ from the plasmid was copied into the NIS found on the BB22OP^Tet^ chromosome and cells in which VPaI-6^Gent^ was only found on the plasmid (**Fig. 2b,c**). Following isolation streaking of a mixed colony, we identified homogenous colonies in which all cells contained a chromosomal copy of VPaI-6^Gent^ inserted into the NIS (**Fig. 2b,c**). These results demonstrate that GMT islands can horizontally transfer between bacteria.

**Fig. 2.**
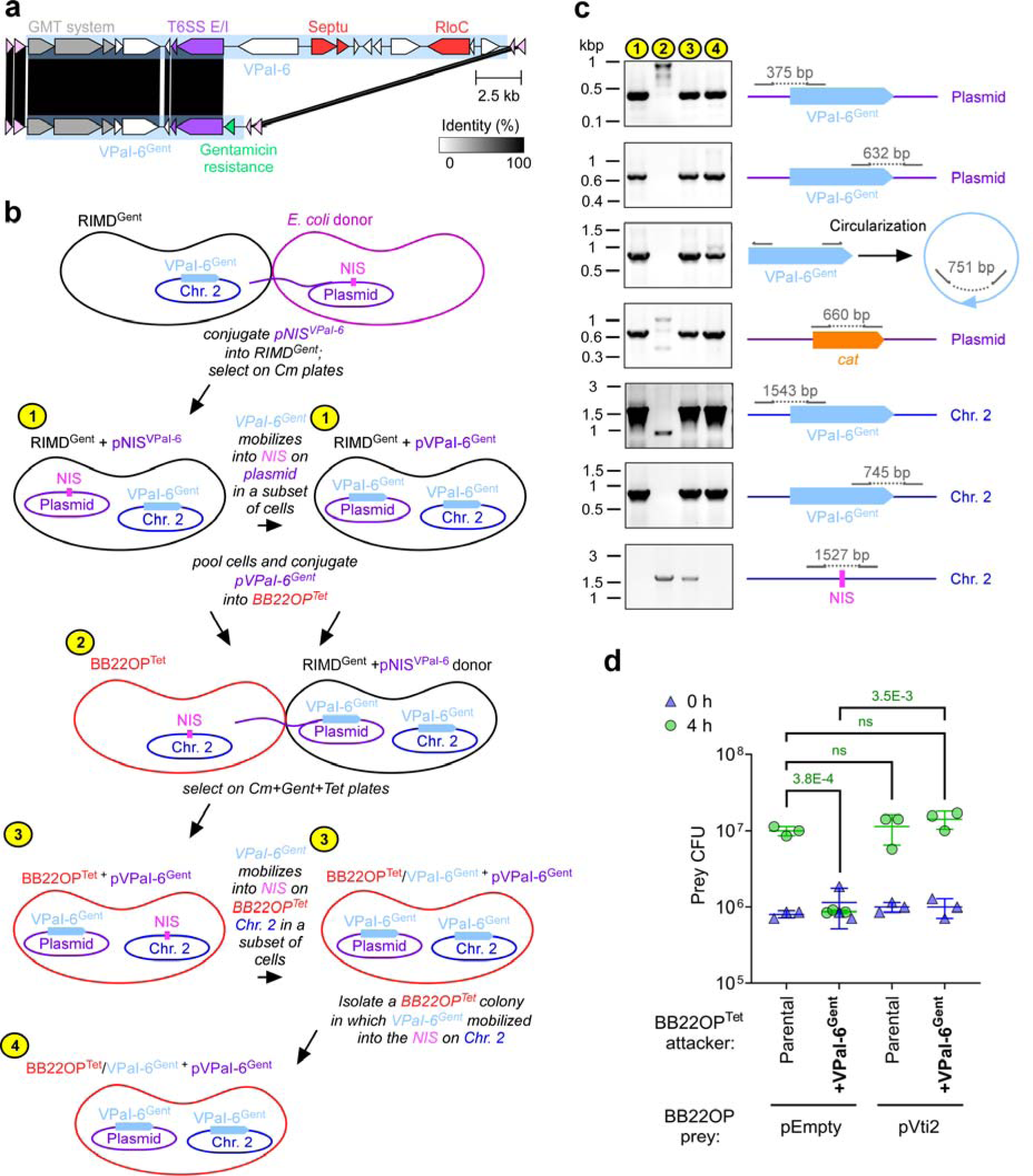
VPaI-6 can be horizontally shared via a conjugatable plasmid. **(a)** Schematic representation of VPaI-6 and VPaI-6^Gent^; gray rectangles denote protein sequence identity percentage. **(b)** An illustration of the assay devised to monitor the plasmid-mediated transfer of VPaI-6^Gent^ between RIMD^Gent^ and BB22OP^Tet^ derivative strains. NIS, naïve insertion site; Chr. 2, chromosome 2; pNIS^VPaI-6^, a plasmid containing a predicted VPaI-6 NIS; Cm, chloramphenicol; Gent, gentamicin; Tet, tetracycline. Numbers in yellow circles denote bacterial populations used for amplicon analysis in (c). **(c)** Agarose gel electrophoresis analysis of the indicated amplicons. The total DNA isolated from the strains denoted by numbers in (b) was used as a template. Samples 1 and 2 are pooled bacteria from both denoted strains in (b). Arrows denote the positions of primers used for each amplicon, and the expected amplicon size is denoted in gray. **(d)** Viability counts (colony forming units; CFU) of the indicated prey strains containing an empty plasmid (pEmpty) or a plasmid expressing the Vti2 immunity protein (pVti2) before (0 h) and after (4 h) co-incubation with the indicated attacker strain. The statistical significance between samples at the 4 h time point was calculated using an unpaired, two-tailed Student’s *t* test; ns, no significant difference (*P* > 0.05); WT, wild-type. Data are shown as the mean ± SD; *n* = 3 independent competition replicates. The data shown are a representative experiment out of at least three independent experiments.

Next, we asked whether the tools within the VPaI-6 cargo can provide a competitive advantage to a bacterium that acquired them. Since VPaI-6^Gent^ contains an antibacterial T6SS effector and immunity pair (VPA1263-Vti2; **Fig. 2a**) absent in the parental BB22OP strain, we hypothesized that after acquiring this island, the T6SS of BB22OP can use the effector to gain competitive advantage. To test this hypothesis, we employed the isolated BB22OP^Tet^ strain containing a chromosomal VPaI-6^Gent^ as an attacker in competition against a parental BB22OP prey strain. As expected, this attacker intoxicated the parental prey compared to a BB22OP^Tet^ attacker strain lacking VPaI-6^Gent^ (**Fig. 2d**). Moreover, expressing the Vti2 immunity protein from a plasmid in the prey strain alleviated this toxicity. These results demonstrate that the BB22OP strain used an antibacterial effector acquired on a GMT island to intoxicate its parental strain, indicating that horizontally shared GMT islands provide a competitive advantage to recipient bacteria.

### GMT islands are rich in defensive and offensive tools

When analyzing the VPaI-6 cargo, we identified a Septu^16^ and a RloC^44^ anti-phage defense systems (VPA1260-VPA1261 and VPA1255, respectively) in addition to the antibacterial T6SS effector and immunity pair (**Fig. 1a**). The co-occurrence of defense systems and a secreted offensive toxin within the same genomic island was not previously reported, prompting us to investigate whether other GMT islands contain a similarly mixed cargo.

We observed considerable variability in the cargo length of the GMT islands for which we identified the borders: between 4.8 and 152.2 kb-long, with a median of ∼13.6 kb (**Fig. 3a**). These lengths differ between bacterial families. Accordingly, the number of genes within GMT islands varies between 3 and 143, with a median of ∼9.6 (**Fig. 3b**). Notably, the cargo of ∼14% of these GMT islands (51 out of 366) comprises only the three genes encoding the GMT system (**Dataset S3**), implying that these GMT proteins are the core components of the MGE.

**Fig. 3.**
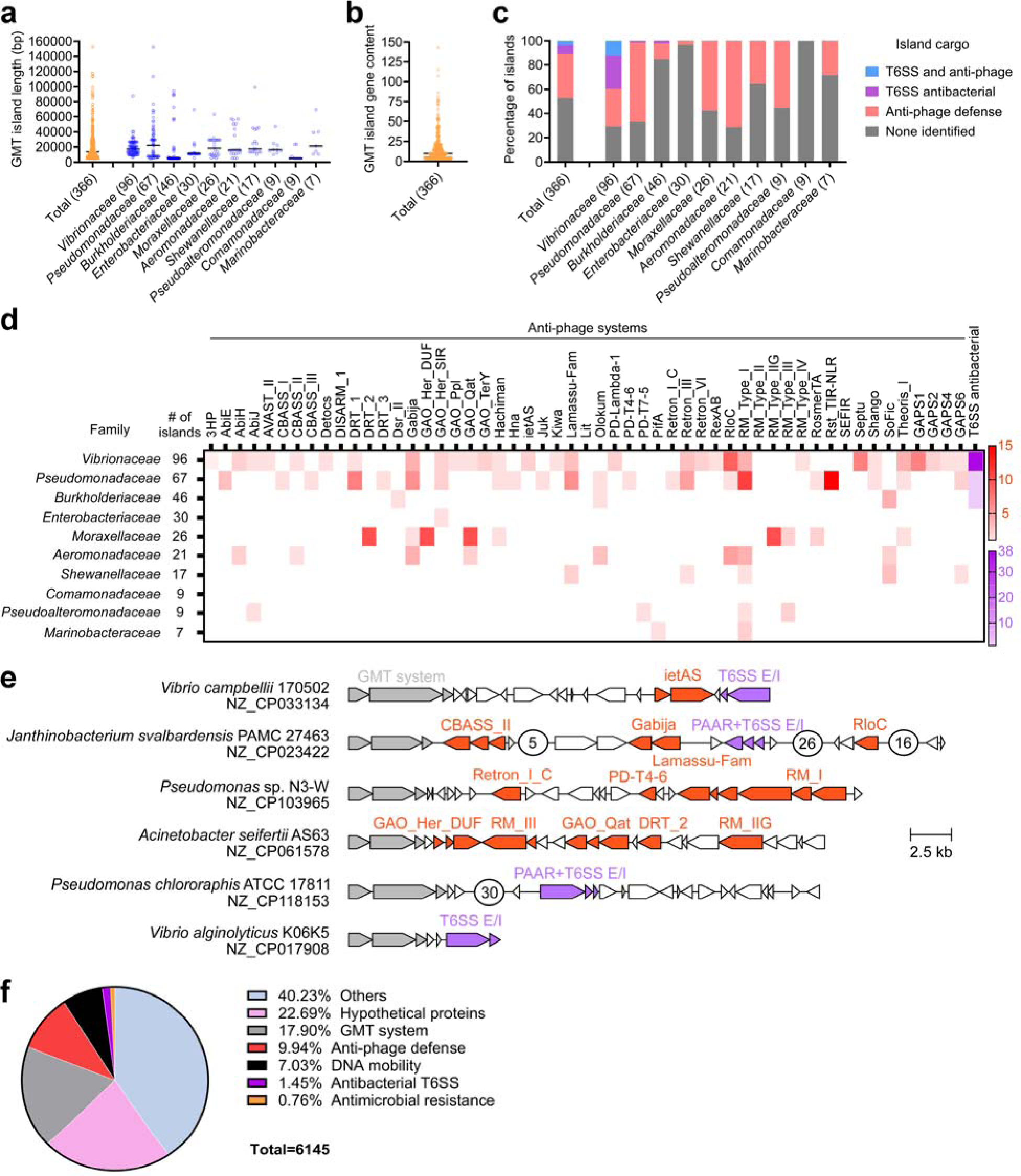
GMT islands contain a diverse cargo of offensive and defensive tools. **(a)** Distribution of GMT island lengths analyzed together (total) or by bacterial family. Black lines denote the median length. **(b)** Distribution of gene number per GMT island. A black line denotes the median gene number. **(c)** Percentage of GMT islands in which we identified an anti-phage defense system, an antibacterial T6SS effector, or both together. **(d)** Distribution of specific anti-phage defense systems and antibacterial T6SS effectors in GMT islands of each bacterial family. Red and purple color gradients denote the number of occurrences, respectively. The analyses in (c-d) include the new anti-phage defense systems identified in this study, as detailed below (i.e., GAPS1, 2, 4, and 6). In (a,c,d), only bacterial families in which we identified the borders of > 5 GMT islands are shown; the number of analyzed islands is denoted in parenthesis next to the family name. **(e)** The gene structure of representative GMT islands with anti-phage defense systems (red), antibacterial T6SS effectors (purple), or both. Encircled numbers denote the number of genes not shown. RefSeq accession numbers are provided. **(f)** A pie chart showing the percentage of GMT island cargo genes associated with the indicated activity or process.

Using domains previously associated with T6SS effectors (i.e., VgrG^1^, PAAR/PAAR-like^45^, MIX^37,38,46,47^, FIX^48^, and Rhs^49^), we identified antibacterial effectors in ∼11% of the examined GMT islands (41 out of 366), all found in genomes harboring a T6SS (**Fig. 3c** and **Dataset S3**). These effectors neighbor a known or predicted immunity gene immediately downstream, and some include a known C-terminal toxic domain (e.g., nuclease or phospholipase). Interestingly, T6SS effectors are prevalent in GMT islands found in members of the *Vibrionaceae* family (∼40%; **Fig. 3c** and **Dataset S3**).

Further analysis of GMT island cargoes using the anti-phage defense system identification servers PADLOC^50^ and DefenseFinder^51^ revealed diverse anti-phage defense systems distributed among most bacterial families (**Fig. 3c-e** and **Dataset S3**). Approximately 40% of the GMT islands contain at least one predicted anti-phage defense system (145 out of 366). Notably, ∼12% of the *Vibrionaceae* GMT islands (12 out of 96) contain both anti-phage defense systems and antibacterial T6SS effectors (**Fig. 3c,e** and **Dataset S3**).

We also found that ∼7% of the analyzed GMT islands (28 out of 366) contain genes associated with antimicrobial resistance, which have been previously reported to reside within MGEs^52–54^ and occasionally also in association with anti-phage defense systems^55^. Notably, no cargo gene encoding a predicted virulence toxin was identified within these 366 GMT islands. Taking the abovementioned results together with a functional classification of GMT island cargo genes (**Fig. 3f**), we propose that GMT islands are akin to armories that stockpile defensive and offensive tools against attacking phages and competing bacteria.

### GMT islands harbor novel anti-phage defense systems

Approximately 22% of the GMT islands’ cargo genes encode proteins annotated as hypothetical, thus not associated with specific processes (**Fig. 3f**). Since anti-phage defense systems are prevalent within GMT islands, and because they often cluster within ‘defense islands’^16,17^, we hypothesized that genes of unknown function within GMT islands encode novel anti-phage defense systems. To test this, we assembled a list of 13 genes and operons found within GMT islands in members of the genus *Vibrio*, which we predicted are novel anti-phage defense systems; we named them GAPS (<u>G</u>MT-encoded <u>A</u>nti-<u>P</u>hage <u>S</u>ystem) 1 to 13 (**Fig. 4a**).

**Fig. 4.**
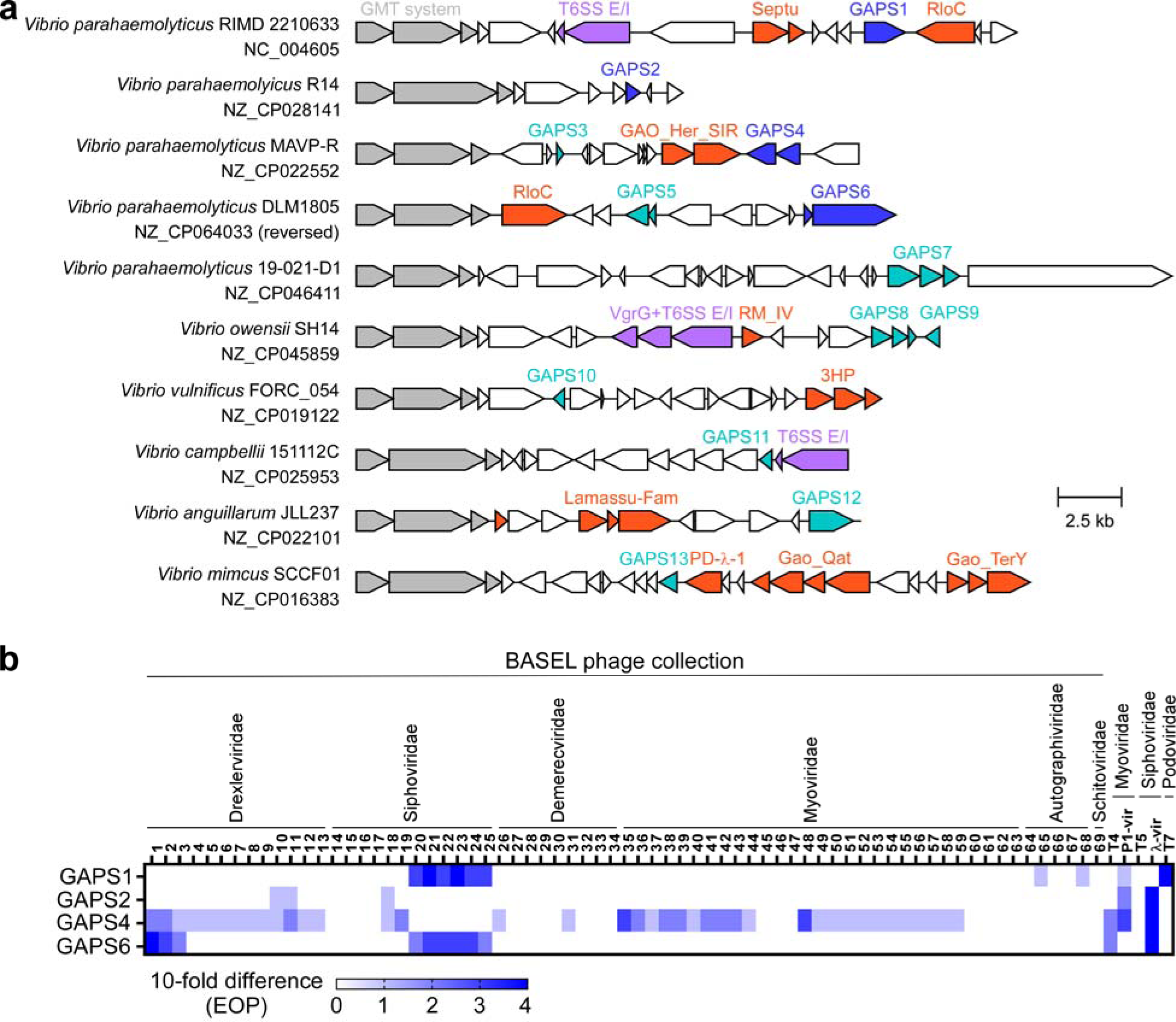
Four new anti-phage defense systems identified within GMT islands. **(a)** The gene structure of GMT islands containing GAPS1-13. GAPSs for which anti-phage activity was experimentally confirmed are denoted in blue; other GAPSs are denoted in turquoise. Known anti-phage defense systems (red) and antibacterial T6SS effectors (purple) are also shown. RefSeq accession numbers are provided. **(b)** The efficiency of plating (EOP), indicating the reduction in plaque numbers determined for *E. coli* expressing the four indicted GAPSs when challenged with 74 coliphages, compared to *E. coli* containing an empty plasmid. Coliphage families are denoted above. The data shown are the average of three independent experiments.

Because a collection of phages that infect a specific *Vibrio* strain is not publicly available, we set out to use *E. coli* as a surrogate host, together with a recently established collection of coliphages^56^, to investigate the role of GAPSs in anti-phage defense. A similar strategy was previously used to identify anti-phage defense systems^57–59^. To this end, we cloned GAPS1-13 into a low-copy number expression plasmid under an arabinose-inducible promoter. *E. coli* strains containing the GAPS-encoding plasmids were challenged with 74 coliphages, comprising the 69 coliphages included in the BASEL phage collection^56^, T7, T4, P1_vir_, T5, and lambda_vir_. We compared the efficiency of plating (EOP) of the different phages in these strains to *E. coli* harboring a control empty plasmid. Remarkably, four candidates: GAPS1 (WP_005477165.1), GAPS2 (WP_174208646.1), GAPS4 (WP_055466293.1 and WP_055466294.1), and GAPS6 (WP_248387294.1 and WP_248387295.1) provided significant protection against various phages, manifested by a reduction of between two and four orders of magnitude in the number of visible plaques that developed on a lawn of bacteria (i.e., the EOP) (**Fig. 4b**). Six GAPSs (GAPS7, GAPS9, GAPS10, GAPS11, GAPS12, and GAPS13) had no significant protective effect against any of the examined phages. Notably, three candidates were considerably toxic to *E. coli* upon expression (GAPS3, GAPS5, and GAPS8); therefore, we could not determine their anti-phage activity. These results support our hypothesis that the cargoes of GMT islands harbor new anti-phage defense systems. Notably, we identified GAPS1, GAPS2, and GAPS6 homologs in several GMT islands (**Fig. 3c,d** and **Dataset S3**)

### GAPS1 belongs to the PD-(D/E)xK superfamily of phosphodiesterases

Three of the newly identified anti-phage defense systems contain predicted domains that may play a role in their activity. GAPS1 and GAPS4 contain a phosphodiesterase domain of the PD-(D/E)xK superfamily^60,61^; the second of the two proteins comprising GAPS6 has a TPR domain and a PINc RNase domain^62^ (**Fig. 5a**; domains were predicted using HHpred^63^). We did not identify similarity to known domains in GAPS2. Prompted by these findings, we further investigated GAPS1, which is encoded within the VPaI-6 GMT island analyzed above.

**Fig. 5.**
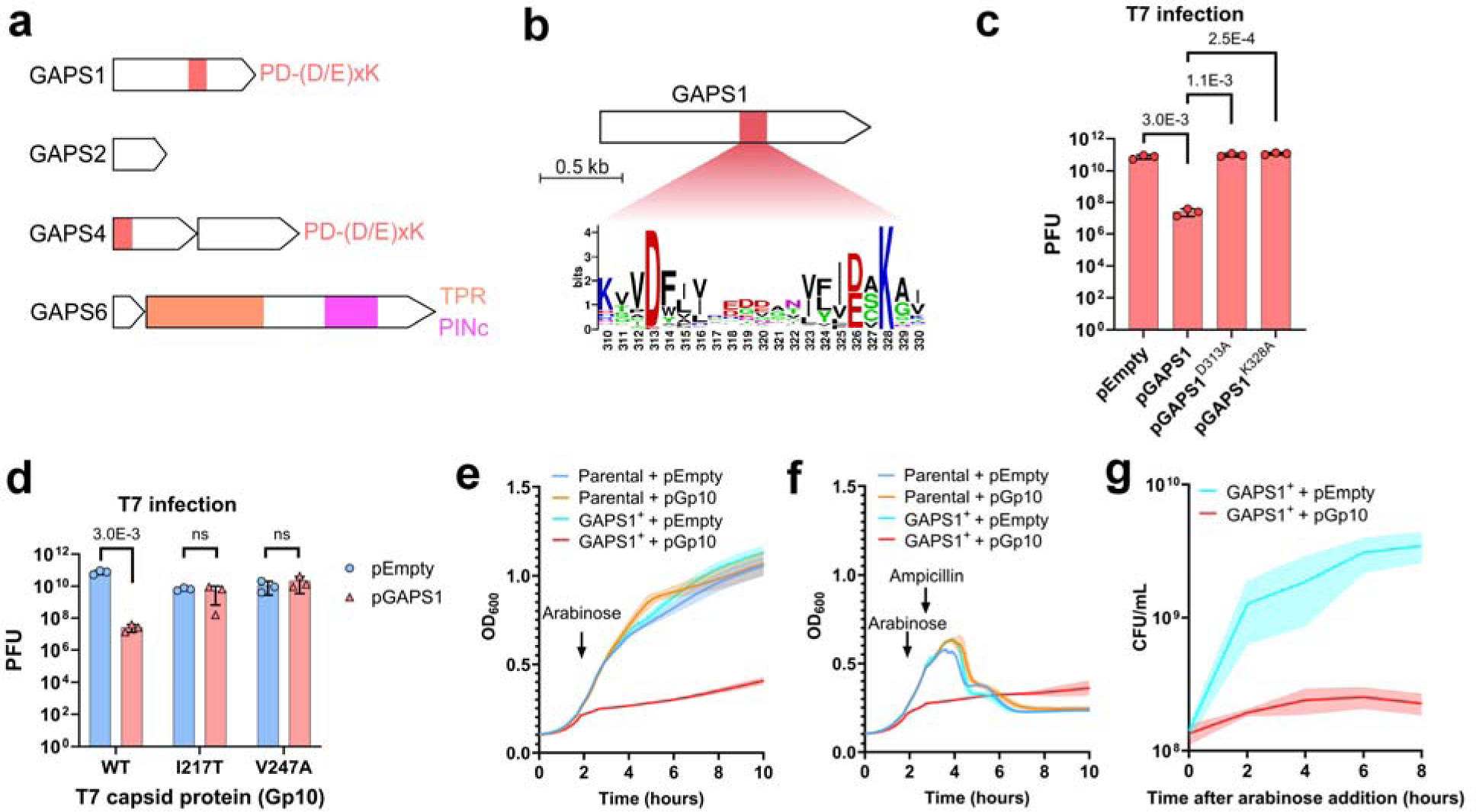
GAPS1 induces cell dormancy upon activation by a phage capsid protein. **(a)** Schematic representation of domains identified in the indicated anti-phage defense systems. **(b)** The conservation logo of the predicted PD-(D/E)xK phosphodiesterase domain active site found in GAPS1 homologs. The residue numbers correspond to the positions in GAPS1 (WP_005477165.1). **(c)** Plaque forming units (PFU) of T7 phage upon infection of *E. coli* strains containing an empty plasmid (pEmpty) or a plasmid for the arabinose-inducible expression of the indicated GAPS1 version. **(d)** PFU of T7 phage, either wild-type (WT) or containing the indicated mutation in the Gp10 capsid protein, upon infection of *E. coli* strains containing an empty plasmid (pEmpty) or a plasmid for the arabinose-inducible expression of GAPS1 (pGAPS1). **(e,f)** Growth of *E. coli* strains in which the chromosomal *ydhQ* gene was deleted (Parental) or replaced with an arabinose-inducible GAPS1 (GAPS1^+^) containing either an empty plasmid (pEmpty) or a plasmid for the arabinose-inducible expression of the T7 phage Gp10 (pGp10). An arrow denotes the time at which arabinose or ampicillin was added. **(g)** Viability (measured as CFU/mL) of GAPS1^+^ *E. coli* strains containing the indicated plasmids after arabinose addition. In (c,d), the data are shown as the mean ± SD of three biological replicates. Statistical significance between samples was determined by an unpaired, two-tailed Student’s *t*-test; ns, no significant difference (*P* > 0.05). In (e-g), a representative experiment out of at least three independent experiments is shown.

GAPS1 is a single protein containing a predicted phosphodiesterase domain toward its C-terminus (**Fig. 5b**). GAPS1 homologs are widespread in Gram-negative and Gram-positive bacteria, and their phylogenetic distribution suggests possible horizontal transfer between bacterial orders (**Extended Data Fig. 5**). They were identified in ∼3.5% of the 294,097 RefSeq genomes analyzed in this study; notably, ∼78% of the identified GAPS1 homologs are encoded in *Klebsiella pneumoniae* genomes (**Dataset S4**). To determine whether the predicted phosphodiesterase domain is required for the anti-phage activity of GAPS1, we substituted D313 and K328 within its conserved PD-(D/E)xK active site with alanines. These substitutions abolished the defensive activity against T7 phage, supporting a role for this domain in GAPS1-mediated anti-phage defense (**Fig. 5c**).

### A phage capsid protein triggers the anti-phage activity of GAPS1

Next, we sought to identify the phage component that triggers GAPS1. One of the phages against which GAPS1 protects is T7 (**Fig. 5c**). We hypothesized that escape mutants of an attacking T7 phage contain mutations in the protein that triggers GAPS1 to avoid system activation. Therefore, we sequenced the genome of four T7 escape phages that formed plaques in the presence of GAPS1. We identified mutations in the capsid protein-encoding gene 10 (Gp10) of all four isolates, including E183A, I217T, and V247A mutations (**Table S1**). Notably, E183, I217, and V247 are spatially close in the folded capsid protein (**Extended Data Fig. 6**).

To determine whether the T7 capsid protein triggers GAPS1-mediated defense, we generated two of the mutations identified above, I217T and V247A, in naïve T7 phages. We validated that additional mutations we identified in the escape isolates are not present in these newly constructed phages, and then tested the ability of GAPS1 to defend against them. We observed no significant difference in the number of mutant T7 phage plaques formed on *E. coli* expressing GAPS1 compared to *E. coli* containing an empty plasmid, indicating that GAPS1 could not defend against the mutant phages (**Fig. 5d**). These findings suggest that the capsid protein triggers GAPS1-mediated defense.

### GAPS1 induces cell dormancy

Anti-phage defense systems protect bacterial populations by inducing cell suicide, often called abortive infection, or by arresting bacterial growth^64–67^. To determine what mechanism is used by GAPS1, we monitored the effect of its activation on bacterial growth and viability in the absence of T7 phage-derived toxins. Expression of the wild-type capsid protein (Gp10) from a plasmid inhibited the growth of *E. coli* expressing a chromosomally inserted GAPS1 (**Fig. 5e**). These arrested cells were protected from ampicillin-induced lysis, indicating that they were not actively dividing or synthesizing peptidoglycan^68^ (**Fig. 5f**). Importantly, we did not observe a reduction in cell viability over time (**Fig. 5g**), demonstrating that Gp10-triggered GAPS1 activity is not bactericidal. These results suggest that GAPS1 is a member of the PD-(D/E)xK phosphodiesterase superfamily that induces cell dormancy rather than cell suicide upon recognizing a phage capsid protein.

### *E. coli* GAPS1 homologs are bona fide anti-phage defense systems

The results demonstrating anti-phage defense against coliphages were obtained by over-expressing an exogenous GAPS1 originating from *V. parahaemolyticus* in *E. coli*. To confirm that *E. coli* GAPS1 homologs protect against coliphages, we cloned three *E. coli* GAPS1 homologs from different strains (EGQ2075554.1, EJP5250929.1, and WP_152927281.1) into the same expression plasmid used to investigate the *Vibrio* GAPS candidates; these homologs share 24-29% amino acid identity with GAPS1 across 64-96% of its length (**Extended Data Fig. 7**). As predicted, these three GAPS1 homologs protected the surrogate *E. coli* against diverse coliphages (**Fig. 6a**).

**Fig. 6.**
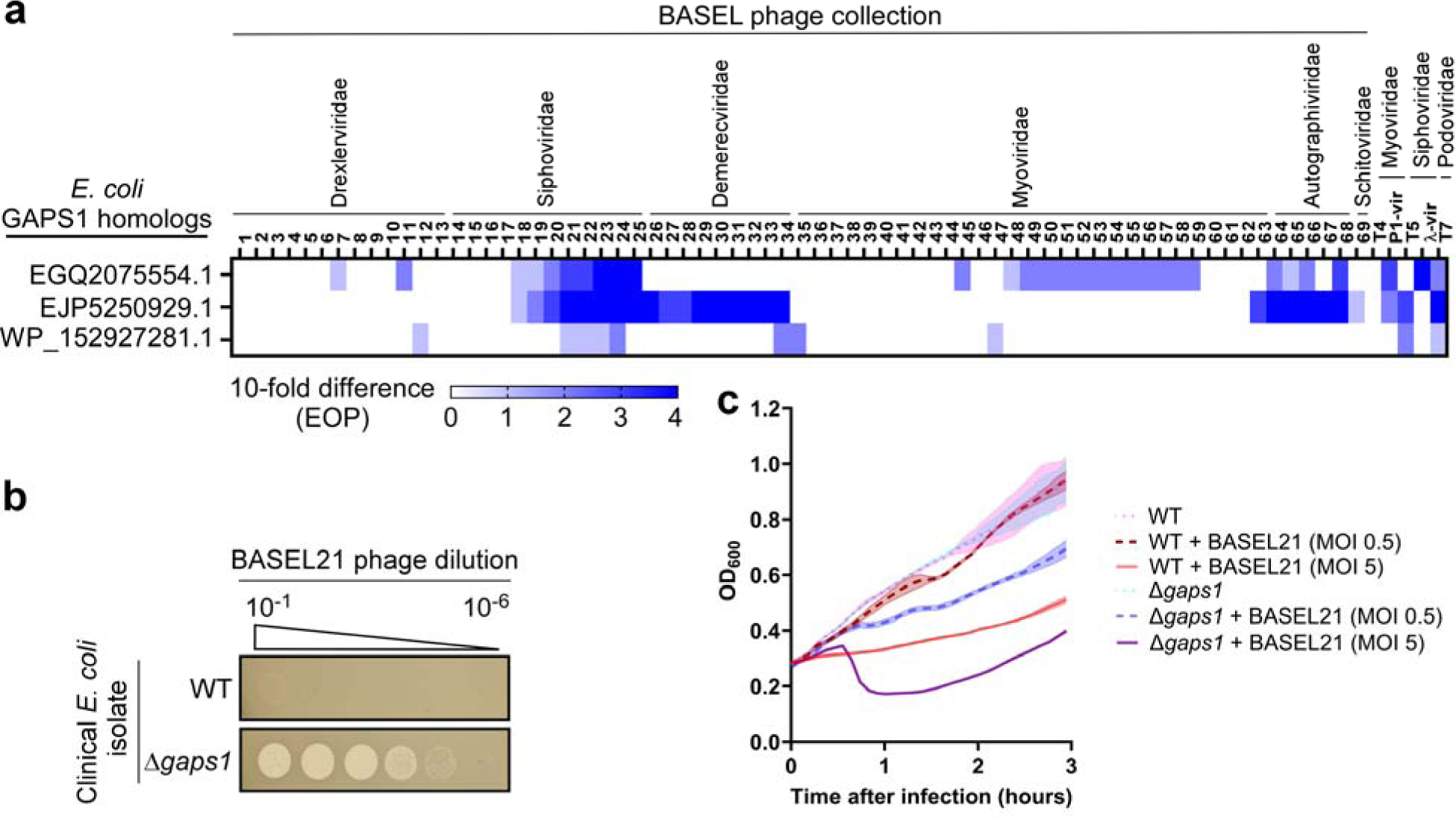
*E. coli* GAPS1 homologs protect against various coliphages. **(a)** The efficiency of plating (EOP) determined for *E. coli* expressing the three indicted GAPS1 homologs when challenged with 74 coliphages, compared to *E. coli* containing an empty plasmid. The data shown are the average of three independent experiments. **(b)** Tenfold serial dilutions of the BASEL collection phage 21 (BASEL21) spotted on lawns of *E. coli* isolate ZH142-A, either wild-type (WT) or with a deletion of its endogenous GAPS1 homolog (Δ*gaps1*). **(c)** Growth of the indicated *E. coli* ZH142-A cells following infection with the BASEL21 phage at MOI of 0.5 or 5. Data are shown as the mean ± SD of three biological replicates. In (b,c), a representative result out of three independent experiments is shown.

Importantly, we obtained a clinical *E. coli* isolate, ZH142-A, naturally encoding a GAPS1 homolog (WP_194242909.1; **Extended Data Fig. 7**). We found that the endogenous GAPS1 homolog is required to protect this strain against predation by BASEL collection phage 21 (BASEL21; **Fig. 6b**). While the growth of the wild-type *E. coli* strain was largely unaffected when challenged with a low phage-to-bacteria ratio (multiplicity of infection [MOI] = 0.5), a high ratio (MOI = 5), in which all bacteria are expected to encounter phage attack, led to growth arrest (**Fig. 6c**). This result is in agreement with the dormancy observed in the surrogate *E. coli* exogenously expressing the *Vibrio* GAPS1 together with the T7 capsid protein (**Fig. 5e**). However, in the absence of the endogenous GAPS1 (Δ*gaps1*), challenging bacteria with a high phage-to-bacteria ratio led to cell lysis, manifested as a drop in the optical density of the bacterial culture, implying successful infection by the phage (**Fig. 6c**). These results confirm that GAPS1 establishes a novel family of anti-phage defense systems.

## Discussion

An evolutionary arms race forces bacteria to acquire new offensive and defensive tools to outcompete rival bacteria and survive phage attacks. Although HGT plays a significant role in this arms race^69^, many mechanisms that mediate HGT in bacteria are poorly understood. Here, we describe GMT, a new system akin to a mobile armory that equips bacteria with defensive and offensive tools.

Anti-phage defense systems have been found in the cargo of MGEs^13,20,28,29^ and shown to often cluster in so-called ‘defense islands’^16,19^. They have also been reported to neighbor other defensive tools, such as antibiotic resistance genes, within MGEs^55,70^. Secreted antibacterial toxins were also identified within MGEs^31–33,71^. However, GMT islands are the first reported example of an MGE carrying defensive and offensive tools together or interchangeably. Although predominantly prevalent in *Vibrionaceae* GMT islands, this phenomenon of mixed offensive and defensive cargo may be common in other types of MGEs found in diverse bacteria. We propose that MGEs previously regarded strictly as ‘defense islands’ should be re-analyzed, considering they might contain new antibacterial offensive tools.

Our results imply that GMT islands are functional mobile elements that employ a replicative mechanism to distribute and insert themselves into specific sites containing inverted repeat sequences (see model in **Extended Data Fig. 8**). Similar replicative mechanisms were previously suggested for transposons^72,73^; however, unlike transposons, the excised and circularized GMT island does not include the flanking repeat sequences, which are probably important for insertion site identification. Even though we found that all three core GMT system proteins are required for the circularization step and, thus, the insertion step, further investigation is required to determine the specific role of each GMT protein in the process.

We propose that plasmids could mediate the dissemination of GMT islands via HGT, as we demonstrated in **Fig. 2**. In support of this notion, we find examples of GMT islands on plasmids encoding conjugation machinery (**Dataset S1**). Some bacteria even contain two identical GMT islands, one on the chromosome and another on a plasmid (**Dataset S2**). In *V. alginolyticus*, for example, the predicted chromosomal and plasmid NISs share inverted repeat sequences with an identical 5’ AAGAGC 3’ core separated by a 14 bp-long spacer (**Extended Data Fig. 9**). Therefore, it is possible that if a NIS is found on a plasmid, a GMT island can replicate itself from the chromosome to the plasmid and then exploit the plasmid to reach other bacteria via HGT.

Many genes within the cargo of GMT islands have no known function. We leveraged the finding that these MGEs are rich in defensive tools to reveal four new anti-phage defense systems. Two of these, GAPS1 and GAPS4, contain predicted domains belonging to the PD-(D/E)xK phosphodiesterase superfamily^61^, which had been previously reported in many anti-phage defense systems^16,20^. Notably, the investigated GAPSs originate from vibrios, yet we examined them in *E. coli* as a surrogate platform against a collection of coliphages. The rationale behind this strategy, which was successfully used by others to identify and investigate anti-phage defense systems^57–59^, is two-fold: (i) a collection of *Vibrio* phages similar to the coliphage BASEL collection^56^ is currently unavailable; (ii) the candidate GAPSs that we investigated originate from different species. To further support the results obtained in the *E. coli* surrogate platform, we showed that an endogenous GAPS1 homolog in a clinical *E. coli* isolate protects the bacterium against a native coliphage (**Fig. 6b-c**). Although we could not confirm their role against phages, the nine GAPSs that did not protect against coliphages may defend against a *Vibrio*-specific phage when expressed in their natural host, or against a phage family that was not included in our coliphage array^74^. Alternatively, additional regulatory or accessory components endogenously found in vibrios might be required for defense activity. In light of our findings, we predict that many other genes within GMT islands encode novel anti-phage defense systems or novel antibacterial toxins.

GAPS1, encoded on VPaI-6, represents a new widespread family of anti-phage defense systems. Being a single protein, GAPS1 probably contains both the sensor that recognizes the phage trigger and the effector domain, which we predict is its PD-(D/E)xK phosphodiesterase domain. By identifying the phage trigger, we could decipher the outcome of GAPS1 activation being cell dormancy rather than cell death. However, since the target of GAPS1 activity remains unknown, it is unclear whether GAPS1 merely inhibits cell growth to halt the progression of the phage infection cycle. It is possible that GAPS1 also actively eliminates the invading phage threat, thus allowing for cell recovery.

Although widespread, the GMT system is predominantly found in gamma-proteobacteria, some beta-proteobacteria, and a handful of Gram-positive families. Nevertheless, our analysis of GMT homologs in this work was conservative, considering only systems highly similar to the trio found in VPaI-6. More distant trios may exist in other bacterial families, and their cargo could contain additional defensive and offensive tools. In future work, we will determine whether GMT systems are regulated, and what is the role of GMT components in the mobility mechanism. We will also decipher how each system identifies a unique and specific insertion site. Further investigation of these intriguing mobile armories will shed light on bacterial interactions, evolution, and HGT.

## Methods

### Strains and media

For a complete list of strains used in this study, see **Table S2**. *Escherichia coli* strains were grown in Lysogeny Broth (LB; 1% [wt/vol] tryptone, 0.5% [wt/vol] yeast extract, and 0.5% [wt/vol] NaCl) or 2xYT (1.6% [wt/vol] tryptone, 1% [wt/vol] yeast extract, and 0.5% [wt/vol] NaCl) at 37°C. *Vibrio parahaemolyticus* strains were grown in Marine Lysogeny Broth (MLB; LB containing 3% [wt/vol] NaCl) and on Marine Minimal Media (MMM) agar plates (1.5% [wt/vol] agar, 2% [wt/vol] NaCl, 0.4% [wt/vol] galactose, 5 mM MgSO_4_, 7 mM K_2_SO_4_, 77 mM K_2_HPO_4_, 35 mM KH_2_PO_4_, and 2 mM NH_4_Cl) at 30°C. Media were supplemented with 1.5% (wt/vol) agar to prepare solid plates. When required, media were supplemented with 35 or 10 µg/mL chloramphenicol (for *E. coli* and V*. parahaemolyticus*, respectively), 50 or 250 µg/mL kanamycin (for *E. coli* and V*. parahaemolyticus*, respectively), or 100 µg/mL ampicillin to maintain plasmids. To induce the expression from P*bad* promoters, 0.04% or 0.2% (wt/vol) L-arabinose was added to the media, as indicated.

### Plasmid construction

Plasmids were constructed with standard molecular biology techniques using the Gibson Assembly method^75^. The Gibson Assembly master mix was obtained from NEB (E2611S). DNA fragments were amplified by PCR from bacterial genomic DNA or from DNA synthesized by TWIST Bioscience, and Gibson Assembly ligations were carried out according to the manufacturer’s instructions. Commercially synthesized DNA, plasmids, and primers that were used in this study are listed in **Table S3**, **Table S4**, and **Table S5**, respectively.

### Constructing *Vibrio parahaemolyticus* mutant strains

For in-frame deletions and gene replacement in *V. parahaemolyticus* RIMD 2210633 or BB22OP, pDM4-based suicide plasmids^76^ were used. Plasmids for gene deletions contained fusions of approximately 600 bp-long sequences upstream and downstream of the region to be deleted in their multiple cloning site (MCS). Plasmids for gene replacement also contained the sequence intended for insertion into the chromosome between the upstream and downstream sequences mentioned above.

To construct the RIMD^VPaI-6_Gent^ strain, wherein the *vpa1254*-*vpa1262* genes were replaced by a gentamicin resistance (Gent^R^) gene downstream of a constitutive promoter, a *cat* promoter amplified from plasmid pBAD33.1^77^ was ligated to the Gent^R^ gene amplified from plasmid pBAD18-Gm. These were then inserted between the sequence upstream of *vpa1253* and the sequence downstream of *vpa1263* in the pDM4 MCS.

To construct the BB22OP^Tet^ strain, wherein the *dns* gene (*vpbb_rs12365*) was replaced by a tetracycline resistance gene (Tet^R^), the Tet^R^ gene (*tetA*) was amplified together with its constitutive promoter tetR/A from *E. coli* IYB5101^78^ and inserted between the *dns* upstream and downstream sequences in the pDM4 MCS.

To construct the RIMD^sfGFP-NIS^ strain, wherein the *vp1393* (*hcp1*) was replaced by a superfolder GFP (sfGFP) harboring a VPaI-6 naïve insertion site, the sequence of sfGFP containing a 9 amino acid-long linker between the 10^th^ and 11^th^ β-strands was amplified from a commercially synthesized plasmid (pTWIST-sfGFP-linker; Twist Biosciences) and inserted between the upstream and downstream sequences of *vp1393* in the MCS of a pDM4 plasmid. Next, the linker sequence was replaced by a 30 bp-long VPaI-6 naïve insertion site.

The described pDM4 constructs were transformed into *E. coli* DH5α (λ-pir) by electroporation, and then conjugated into *V. parahaemolyticus* via tri-parental mating with the help of an *E. coli* conjugation helper strain. Next, trans-conjugants were selected on MMM agar plates containing 10 µg/mL chloramphenicol or, when necessary, supplemented with 5 µg/mL tetracycline or 25 µg/mL gentamicin. The resulting trans-conjugants were grown on MMM agar plates containing 15% [wt/vol] sucrose for counter-selection and loss of the *sacB*-containing pDM4. Deletions were confirmed by PCR.

### GMT island mobility assays

#### Transfer from a V. parahaemolyticus chromosome to a plasmid

pBAD33.1 plasmids, either empty or containing a 30 bp-long VPaI-6 naïve insertion site (pNIS^VPaI-6^) or its mutated forms, or containing a naïve insertion site for the GMT island found in *V. parahaemolyticus* 04.2548 (pNIS^04.2548^) were transformed into *E. coli* Neb5α and subsequently introduced into the indicated *V. parahaemolyticus* strains via tri-parental mating. The resulting conjugated colonies were selectively grown on MMM agar plates supplemented with 10 µg/mL chloramphenicol. Then, all the colonies that grew on the selective plate were harvested, resuspended in LB media, and subjected to total genomic DNA isolation using the PrestoTM Mini gDNA isolation kit. To identify instances in which the GMT island of interest mobilized from the chromosome into the pBAD33.1-based plasmid, 110 ng of isolated total DNA was used as template to perform PCR using primer sets intended to amplify: (i) a fusion between the plasmid and the 5’ end of the GMT island, (ii) a fusion between the plasmid and the 3’ end of the GMT island, (iii) a fusion between the 5’ and 3’ ends of the GMT island (i.e., circularization), and (iv) a chloramphenicol resistance gene (*cat*; Cm^R^) found in the pBAD33.1 backbone (used as an internal control for plasmid presence). PCR products were resolved on a 0.8% agarose gel and visualized with EtBr staining.

#### Transfer between V. parahaemolyticus strains

To monitor the transfer of VPaI-6^Gent^ between RIMD^Gent^ and BB22OP^Tet^, a colony of RIMD^Gent^ in which the mobilization of VPaI-6^Gent^ to the pNIS^VPaI-6^ plasmid was confirmed via PCR amplifications (as described above) was used as a donor in tri-parental mating together with an *E. coli* conjugation helper strain and BB22OP^Tet^ recipient cells. The resulting trans-conjugates were grown on MMM agar plates supplemented with 10 µg/mL chloramphenicol, 5 µg/mL tetracycline, and 25 µg/mL gentamicin to select for BB22OP^Tet^ colonies containing a pNIS^VPaI-6^ plasmid with a VPaI-6^Gent^. The transfer was confirmed using PCR amplifications.

#### Transfer from a plasmid to a naïve insertion site in the V. parahaemolyticus BB22OP chromosome

To monitor the transfer of VPaI-6^Gent^ from a plasmid to the natural VPaI-6 naïve insertion site found on chromosome 2 of *V. parahaemolyticus* BB22OP, a single colony of BB22OP^Tet^ containing a pNIS^VPaI-6^ plasmid with a VPaI-6^Gent^ (described in the previous section) was re-streaked on a selective plate to isolate colonies in which VPaI-6^Gent^ was found in the bacterial chromosome. The transfer was confirmed using PCR amplifications.

#### Discriminating between a replicative and conservative transfer mechanism

To determine whether VPaI-6 mobilizes via a replicative (copy-and-paste) or conservative (cut-and-paste) mechanism, a fluorescent RIMD^sfGFP-NIS^ colony was streaked on a plate and incubated for 16 hours at 30°C. The colonies were then inspected under blue light to identify a colony that lost the GFP fluorescence, indicative of inactivation of the sfGFP open reading frame, likely by insertion of VPaI-6 into the intragenic naïve insertion site. This colony was then re-streaked, and a single colony was used to extract genomic DNA and determine the location of VPaI-6 via PCR amplifications.

### Bacterial competition assays

The indicated attacker and prey *V. parahaemolyticus* BB22OP strains were cultured overnight in MLB with appropriate antibiotics, normalized to an OD_600_ of 0.5, and then mixed at a 4:1 (attacker:prey) ratio in triplicate. Subsequently, 25 μL of the mixtures were spotted onto MLB agar competition plates and incubated at 30°C for 4 hours. To determine the colony-forming units (CFU) of the prey strains at tLJ=LJ0LJhours, 10-fold serial dilutions were plated on MMM agar plates supplemented with 10 µg/mL chloramphenicol and 250 µg/mL kanamycin. After 4 hours of co-incubation of the attacker and prey mixtures on the competition plates, the bacteria were harvested, and the CFUs of the surviving prey strains were determined as described above. Prey strains harbored a pVSV209^79^ plasmid for selective growth. A representative result out of three independent experiments is shown.

### Plaque assays

The phages used in this study are listed in **Table S6**. Phages were propagated on *E. coli* K12 MG1655 ΔRM. To determine the effect of the 13 putative defense system (GAPS1-13) against coliphages T4, T5, T7, lambda_vir_, P1_vir_, and the 69 phages included in the BASEL collection^56^, *E. coli* K12 MG1655 ΔRM strains harboring the indicated pBAD33.1-based plasmids were grown overnight in LB supplemented with chloramphenicol and 0.2% (wt/vol) D-glucose (to repress expression from the P*bad* promoter) at 37°C. Overnight cultures were washed twice to remove any remaining glucose, and then 350 μL of each culture were mixed with 7 mL of 0.7% (wt/vol) molten agar supplemented with 0.2% (wt/vol) L-arabinose, 10 mM MgSO_4_, and 5 mM CaCl_2_. The mixture was poured onto a 1.5% (wt/vol) agar plate supplemented with chloramphenicol and 0.2% (wt/vol) L-arabinose, and the plates were left to dry. Tenfold serial dilutions of all the phages were prepared, and 7.5 μL of each dilution were spotted on the dried plates. The plates were incubated overnight at 37°C. The following day, the plaques were counted and the plaque forming units (PFU/mL) were calculated. For dilution spots in which no individual plaques were visible but a faint zone of lysis was observed, the dilution was considered as having ten plaques, as previously described^80^. Plaque assays with *E. coli K12* MG1655 ΔRM containing plasmids for the expression of GAPS1 mutants and homologs were performed similarly. This protocol was also used to investigate the ability of BASEL collection phage 21 to form plaques on *E. coli* ZH142-A and its Δ*gaps1* mutant, except the plates did not include L-arabinose or antibiotics.

### Isolation of T7 escape phages

*E. coli* BW25113 harboring an empty pBAD33.1 or one encoding GAPS1 (pBAD33.1-GAPS1) was grown overnight in LB supplemented with chloramphenicol and 0.2% (wt/vol) D-glucose at 37°C. The cells were washed twice and mixed with 0.7% (wt/vol) molten agar supplemented with 0.2% (wt/vol) L-arabinose, and poured onto a 1.5% (wt/vol) agar plate supplemented with 0.2% (wt/vol) L-arabinose. After the agar dried, tenfold serial dilutions of a T7 phage suspension were spotted onto the plate, and the plate was incubated overnight at 37°C. Individual plaques growing at the highest dilution on the plates containing *E. coli* expressing GAPS1 were isolated and propagated on naïve *E. coli* BW25113 cells harboring pBAD33.1-GAPS1 in LB supplemented with chloramphenicol and 0.2% (wt/vol) L-arabinose at 37°C. The escape phages were confirmed by spotting on plates with *E. coli* expressing GAPS1.

For phage genomic DNA isolation, high titers of escape phages and a parental wild-type phage were prepared (∼1.0E^11^ PFU/mL). Approximately 40 mL of lysates of each escape phage were mixed with 10% (wt/vol) PEG 8000 and 3 M NaCl, and incubated overnight at 4°C. The lysate-PEG mixture was then centrifuged at 10000 x *g* for 15 minutes to collect the phage pellet. The pellet was re-suspended using resuspension buffer from the Presto^TM^ Mini gDNA isolation kit, and the phage genomic DNA was isolated following the manufacturer’s protocol.

Illumina whole-genome sequencing was carried out at SeqCenter (Pittsburgh, PA, USA; https://www.seqcenter.com/). Sample libraries were prepared using the Illumina DNA Prep kit and IDT 10 bp UDI indices, and sequenced on an Illumina NextSeq 2000, producing 2 x 151 bp reads. Mutations were identified using variant calling (SeqCenter). Only mutations that were found in the escape mutant genomes and that were not in the sequenced parental T7 phage are reported in **Table S1**.

### Constructing T7 phage mutants

T7 mutants were constructed using the pORTPHAGE method^81^, a MAGE^82,83^-based system for the mutagenesis of bacteriophages. Briefly, *E. coli* K-12 strain harboring the pORTMAGE-Ec1 plasmid was grown to reach early log phase (OD_600_ ∼0.3). Then, 1 mM m-toluic acid was added to induce the expression of recombineering proteins. The cells were made electrocompetent and then transformed with mutating oligonucleotides. After electroporation, the culture was infected with a wild-type T7 phage and incubated until complete lysis occurred. The final lysate was cleared using chloroform, diluted, and then plated using the soft agar overlay method to screen for individual mutated plaques. Single plaques were picked, suspended in LB, and used as templates for PCR amplification and sequencing to identify mutants.

### Chromosomal integration of GAPS1

GAPS1 was introduced into the chromosome of *E. coli* BW25113 in place of *ydhQ* (*ydhQ*::GAPS1) using the red recombination system, as previously described^84^. Briefly, *E. coli* BW25113 cells harboring pSim6 were grown overnight in LB supplemented with ampicillin at 30°C. Overnight cultures were diluted 1:100 in 35 mL of fresh LB supplemented with ampicillin and grown to an OD_600_ of ∼0.5. The red recombinase system was then heat-induced for 20 minutes in a shaking water bath at 42°C. Immediately after induction, the cells were chilled on ice and pelleted by centrifugation. The cell pellets were washed thrice with ice-cold water and resuspended in 200 μL of ice-cold water. The gene encoding GAPS1 under P*bad* promoter control, along with a kanamycin-resistance cassette, was amplified together with flanking sequences identical to flanking sequences 50 bp upstream and downstream of the chromosomal *ydhQ*. The amplified DNA was treated with DpnI, and then run on an agarose gel and purified. The purified DNA was electroporated into *E. coli*, and bacteria were allowed to recover in 2xYT broth supplemented with 0.2% (wt/vol) D-glucose for two hours at 30°C. The transformed cells were then plated onto a 1.5% (wt/vol) agar plate supplemented with 25 μg/mL kanamycin. The integration of GAPS1 was verified by PCR. Bacteria were cured of the pSIM6 plasmid, and the recombinant cells were electroporated with pCP20 plasmid. The kanamycin cassette was flipped out by inducing the pCP20 plasmid at 42°C.

### Deleting the GAPS1 homolog in *E. coli* ZH142-A

The GAPS1 homolog was deleted from the chromosome of *E. coli* ZH142-A using the lambda red recombination system, and replaced with a kanamycin resistance cassette. A single colony of bacteria containing the pSim6 plasmid was grown overnight in LB supplemented with ampicillin at 30°C. The overnight culture was diluted 1:100 in 35 mL of fresh LB supplemented with ampicillin and grown to an OD_600_ of ∼0.5. The red recombinase system was then heat-induced for 20 minutes in a shaking water bath at 42°C. Immediately after induction, the cells were chilled on ice and pelleted by centrifugation. The cell pellets were washed thrice with ice-cold water and resuspended in 200 μL of ice-cold water. Then, these cells were electroporated with the following DNA: The kanamycin-resistance cassette, along with the flippase recognition target (FRT) sites, was amplified from *E. coli* BW25113ΔydhQ::kan using primer pairs with overhang sequences homologous to the 50 bp of the 5’ and 3’ sequences flanking the GAPS1 homolog. The amplified DNA was treated with DpnI restriction enzyme, run on an agarose gel, and purified.

Following electroporation, the cells were allowed to recover in 2xYT broth for two hours at 30°C. The transformed cells were then plated onto a 1.5% (wt/vol) agar plate supplemented with 25 μg/mL kanamycin. The replacement of the GAPS1 homolog with the kanamycin resistance cassette was verified by PCR. Bacteria were subsequently cured of the pSIM6 plasmid^85^.

### *E. coli* toxicity and viability assays

*E. coli* BW25133 Δ*ydhQ* and *E. coli* BW25133 *ydhQ*::GAPS1 cells harboring an empty plasmid or a plasmid for the arabinose-inducible expression of the T7 phage gene10 (encoding the Gp10 capsid protein) were grown overnight at 37°C in LB supplemented with kanamycin and 0.2% (wt/vol) D-glucose. Overnight cultures were diluted to an OD_600_ = 0.02 in fresh LB supplemented with kanamycin, and 200 µL were transferred into 96-well plates in triplicate. Cells were grown under continuous shaking (205 RPM) in a Tecan Infinite M Plex plate reader at 37°C. After two hours, the expression of Gp10 and GAPS1 was induced by adding L-arabinose to a final concentration of 0.2% (wt/vol). OD_600_ readings were acquired every 10 minutes. A similar procedure was used to determine the effect of adding ampicillin (100 μg/mL) 1 hour after arabinose addition.

To determine cell viability after induction, bacteria were collected at the indicated time points after arabinose addition. Tenfold serial dilutions of each culture were spotted on agar plates supplemented with kanamycin and 0.2% (wt/vol) D-glucose (to repress arabinose-induced expression). The plates were incubated overnight at 37°C, and the CFU/mL of each culture were determined the following day.

To monitor bacterial growth upon infection with BASEL collection phage 21, *E. coli* ZH142-A wild-type and Δ*gaps1* mutant strains were grown overnight at 37°C in LB. Overnight cultures were diluted 1:100 in 10 mL of fresh LB and grown to an OD_600_ of ∼0.3. The phage was then added to the bacterial cultures at the indicated MOI, and 200 µL of cells were transferred into 96-well plates in triplicate. Cells were grown under continuous shaking (205 RPM) in a Tecan Infinite M Plex plate reader at 37°C. OD_600_ readings were acquired every 10 minutes.

### Identification of GMT islands

GMT islands were identified by performing the following steps.

#### Construction of position-specific scoring matrices (PSSMs) of GMT proteins

The PSSMs of VPA1270 (GmtY), VPA1269 (GmtZ), and VPA1268 (GmtX) were constructed using full-length sequences from *Vibrio parahaemolyticus* RIMD 2210633 (WP_005477115.1, WP_005477239.1, and WP_005477284.1, respectively). To improve the identification of GMT proteins, additional PSSMs of VPA1269 and VPA1268 were constructed using full-length sequences from *Vibrio parahaemolyticus* R14 (WP_108745444.1 and WP_085344822.1, respectively). Online PSI-BLAST (https://blast.ncbi.nlm.nih.gov) was employed to construct all PSSMs. In each case, five iterations of PSI-BLAST against the RefSeq protein database were performed. A maximum of 500 hits with an expect value threshold of 10^−6^ and a query coverage of 70% were used in each iteration of PSI-BLAST. Files containing PSSM information were downloaded from the website and were used later in RPS-BLAST analysis (see below).

#### Identification of bacterial genomes containing GMT systems

A local database containing the RefSeq bacterial nucleotide and protein sequences was generated (last updated on August 21, 2023). RPS-BLAST was used to identify GmtY homologs in the local database. The results were filtered using an expect value threshold of 10^−6^ and a query coverage of 70%. Analysis was limited to complete genomes (NCBI assembly level: complete genome or chromosome). Subsequently, the genomic neighborhood of GmtY-containing genomes was analyzed as described before^47,86,87^. The results were further analyzed to identify bacterial sequences containing the three GMT proteins in tandem. Cases where an unrelated protein was inserted between GMT proteins (*e.g.*, a transposase) were accommodated. A list of GMT proteins and adjacently encoded proteins is provided in **Dataset S1.**

#### Identification of closely related genomes

First, the sequences of *rpoB*, coding for DNA-directed RNA polymerase subunit beta, were retrieved from the local database for all RefSeq bacterial genomes. Partial and pseudo-gene sequences were excluded. A nucleotide database of *rpoB* genes was generated. Next, BLASTN was performed using the sequences of *rpoB* from the GMT-containing genomes as queries to identify *rpoB* homologs with high sequence identity (at least 90% over at least 90% of the sequence). The BLASTN results were analyzed and a list of closely related genomes was generated for each GMT-containing genome.

#### Identification of genomic accessions in closely related genomes that are homologous to sequences flanking GMT systems

The nucleotide sequences of the GMT systems and their 5’ and 3’ flanking regions, up to 200 kbp of either side of *GmtY*, were retrieved. These sequences were used as query in BLASTN against the nucleotide sequences of closely related genomes. The results were filtered to include local alignments that are of ≥1 kbp length with ≥80% identity between aligned sequences. The alignments were further analyzed to identify separate alignments belonging to the same genomic accessions that flank the GMT systems but do not include them (**Fig. S8a**). The alignments were required to be with the same strand of the subject accession. The distances between the positions of the alignments in the subject accessions were required to be ≤100 bp (**Fig. S8a**). In addition to the above criteria, the sequences upstream and downstream to the GMT islands were required to contain sequence alignments to the subject accessions in at least 4 kbp out of 10 kbp upstream and downstream sequences. The aim of this step was to remove false alignments due to frequent sequences (e.g., transposases) (**Fig. S8b**).

#### Identification of GMT Island borders

The alignments meeting all the abovementioned requirements were grouped together to determine the 5’ and 3’ borders of GMT islands (**Figure S8c**). First, the consensus values of the borders were deduced based on the most frequently occurring values. Then, the putative borders were ranked based on the following criteria: (i) distance between subject alignments ≤20 bp, (ii) upstream and downstream alignments ≥5 kbp, (iii) borders are ±10 bp from consensus values, and (iv) borders are exactly the same as the consensus values. The putative borders with the highest ranking were selected for further analysis (**Dataset S2**).

#### Analysis of the putative entry sites

The predicted entry site for each GMT island was determined according to positions of the alignments in the subject accessions. Entry sites were defined as ‘intragenic’ or ‘intergenic’ if located inside or outside genes, respectively (**Fig. S8d**). The sequences located 25 bp from the ends of the predicted entry sites were analyzed to identify direct and inverted repeats (**Fig. S8e**). Briefly, to identify direct repeats, all possible sub-sequences located in the first sequence were searched in the second sequence. To identify inverted repeats, the search was performed in the reverse complement. The minimal repeat size was set to 5 nucleotides, and the longest identified repeats were saved.

### Analysis of GMT island cargoes

T6SS effectors were identified by the presence of T6SS effector-specific domains (i.e., MIX^37,47^, FIX^48^, Rhs^49^, PAAR and PAAR-like^45^, Hcp^1^, and VgrG^1^), determined by NCBI Conserved Domain Database (CDD)^39^ (see below) or using previously-constructed PSSMs. Predicted toxic domains of T6SS effectors were identified using CDD or by similarity detection using hidden Markov modeling (HHpred^63^). Small genes downstream of T6SS effectors were annotated as putative immunity genes.

Anti-phage defense systems were identified using the PADLOC^50^ and DefenseFinder^51^ tools. In the case of PADLOC, amino acid sequences and gff3 files of the complete genomes were provided as input. In the case of DefenseFinder, amino acids sequences, ordered according to their position in the genomes, were provided as input. The anti-phage defense systems described in this work were identified by constructing PSSMs for proteins belonging to the systems and identification of homologs using RPS-BLAST. PSSMs of GAPS1, GAPS2, GAPS4a, GAPS4b, GAPS6a, and GAPS6b were constructed using full-length sequences (WP_005477165.1, WP_174208646.1, WP_055466293.1, WP_055466294.1, WP_248387294.1, and WP_248387295.1, respectively). PSI-BLAST was performed as described above for the GMT system. RPS-BLAST results were filtered using an expect value threshold of 10^−15^ and a minimal coverage of 70%. With regard to GAPS4 and GAPS6, all proteins belonging to these systems were required for the systems to be counted.

DNA mobility elements were identified using blast search in the mobileOG database (Beatrix 1.6 v1^88^) and by a manual search for protein descriptions containing ‘transposase’, ‘recombinase’, ‘conjugation’, or ‘integrase’ keywords. Antimicrobial resistance genes were identified using a blast search in the NCBI Pathogen Detection Reference Gene Catalog, available from The NCBI Pathogen Detection Project [Internet]. Bethesda (MD): National Library of Medicine (US), National Center for Biotechnology Information. 2016 May [downloaded: 2024 May 13]. Available from: https://www.ncbi.nlm.nih.gov/pathogens/. Virulence toxins were identified using blast searches in the Virulence Factor Database (VfDB^89^) and in Bastion-HUB database^90^. Blast results from searches in the various databases were manually assessed, and genes encoding transcription regulators were excluded. Partial and pseudo-genes were not included in the analysis.

### Identification of conserved domains

The CDD and related information were downloaded from NCBI on August 27, 2023^39^. RPS-BLAST was employed to identify conserved domains in protein sequences and the output was processed using the Post-RPS-BLAST Processing Utility v0.1. The expect value threshold was set to 10^−5^.

### Construction of phylogenetic trees

Phylogenetic analysis of bacterial strains was conducted using the MAFFT server (mafft.cbrc.jp/alignment/server/)^91^. The nucleotide sequences of *rpoB* were aligned using MAFFT version 7 (FFT-NS-i)^92^. Partial and pseudo-gene sequences were not included in the analysis. The evolutionary history was inferred using the neighbor-joining method^93^ with the Jukes-Cantor substitution model (JC69). The indicated evolutionary distances are in the units of the number of base substitutions per site.

The phylogenetic tree of GmtY and GAPS1 were constructed by performing the following steps. First, protein sequences were aligned using CLUSTAL Omega^94^. Then, evolutionary analyses were conducted in MEGA X^95^. In the case of GmtY, the evolutionary history was inferred by using the Maximum Likelihood method and the LG+G+I model^96^. In the case of GAPS1, the Maximum Likelihood method and the LG+G+I+F model were used. Both models were found to have the lowest BIC (Bayesian Information Criterion) scores among 56 different amino acid substitution models that were analyzed in MEGA X. The analysis of GmtY involved 366 amino acid sequences and 375 conserved sites. The analysis of GAPS1 involved 833 amino acid sequences and 264 conserved sites. The trees were visualized using iTOL^97^ (https://itol.embl.de/).

### Illustration of conserved residues using Weblogo

The protein sequences of GAPS1 homologs were aligned using CLUSTAL Omega^94^. Aligned columns not found in representative proteins were discarded. The conserved residues were illustrated using the WebLogo server (weblogo.berkeley.edu)^98^.

### Multiple sequence alignment of *E. coli* GAPS1 homologs

The amino acid sequences of EGQ2075554.1, EJP5250929.1, WP_152927281.1, WP_194242909.1, and GAPS1 (WP_005477165.1) were aligned using Clustal W^99^ in MEGA X^95^. Similarity and identity shading was done in ESPript 3.0^100^.

## Supporting information

Supplementary Information (Extended Data Figures, Supplementary file captions, and Supplementary References

## Acknowledgments

We thank members of the Salomon, Qimron, and Bosis laboratories for helpful discussions and suggestions. We also thank Andrea Endimiani (University of Bern) for gifting us the *E. coli* ZH1420A strain, and Alexander Harms (ETH Zurich) for generously sharing the BASEL phage collection. DS and EB received funding from the Israel Science Foundation (ISF grant number 1362/21). UQ is supported by the European Research Council – Horizon 2020 research and innovation program, grant no. 818878. UQ has also received funding from the Israeli Ministry of Health in the framework of the ERANET-JPI-AMR, grant no. 15370. KK was supported by a PhD Scholarship from the Tel Aviv University Center for Combatting Pandemics.

