## Supplementary Information (Extended Data Figures, Supplementary file captions, and Supplementary References for "Gamma-Mobile-Trio systems define a new class of mobile elements rich in bacterial defensive and offensive tools"

Extended Data Figures 1-10

Supplementary Tables S1-S6 (captions)

Supplementary Datasets S1-S4 (captions)

Supplementary Files S1-S2 (captions)

Supplementary References

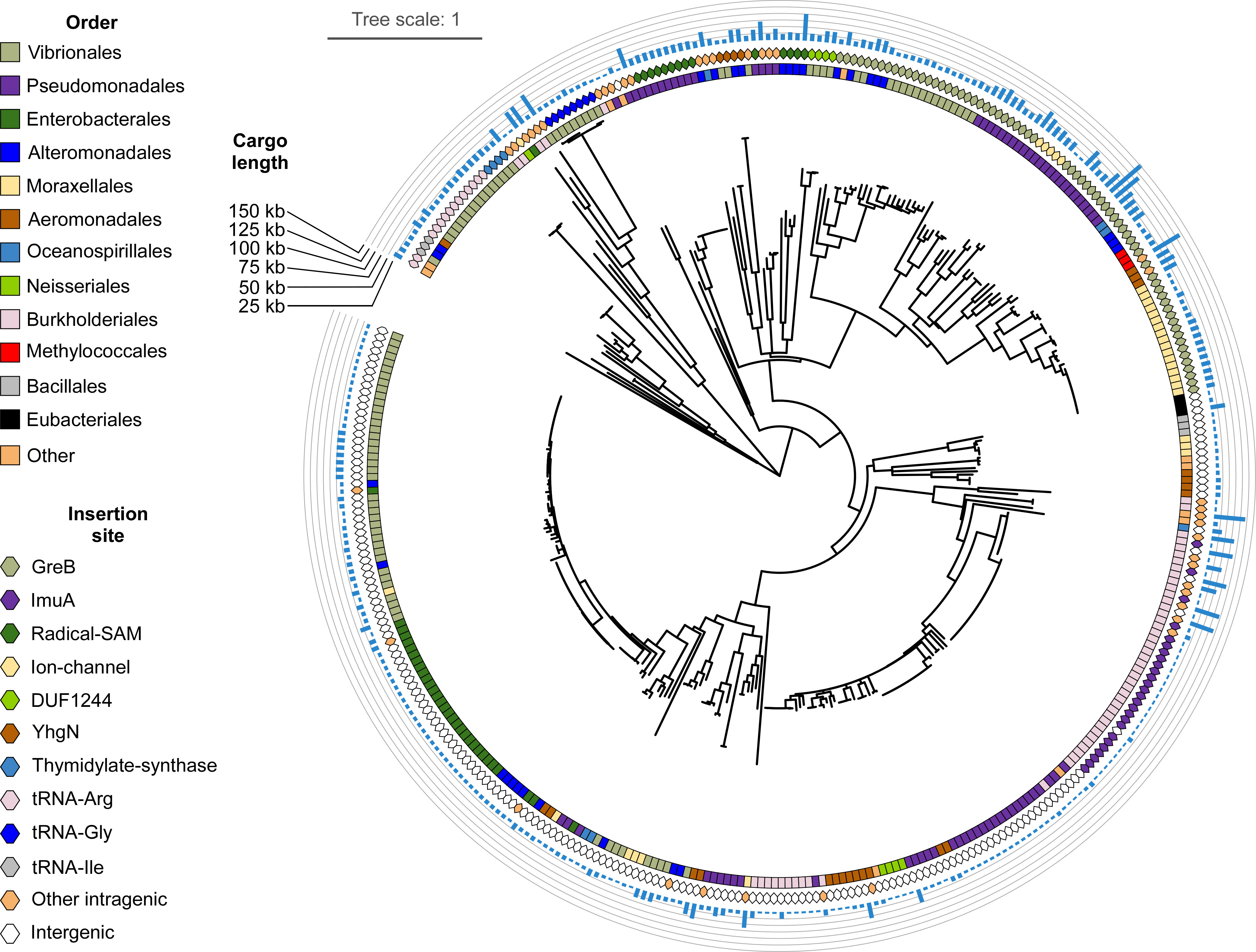
**Extended Data Figures**

**Extended Data Fig. 1. Closely related GmtY proteins are found in diverse bacterial orders.** Phylogenetic distribution of GmtY proteins encoded within GMT islands for which an insertion site was identified. The bacterial order and the type of predicted GMT island insertion site are denoted. Blue bars denote the GMT island cargo length. The evolutionary history was inferred by using the Maximum Likelihood method and LG+G+I model. The tree is drawn to scale, with branch lengths measured in the number of substitutions per site. Evolutionary analyses were conducted in MEGA X.

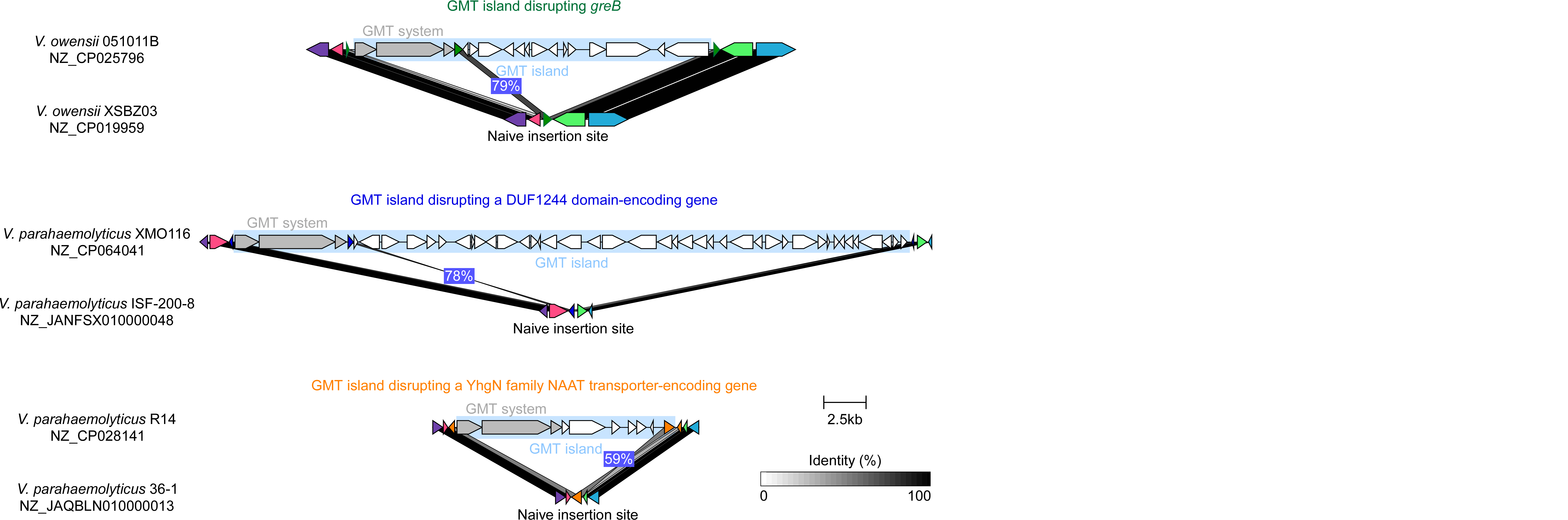

**Extended Data Fig. 2. GMT islands with a predicted intragenic insertion site contain a homolog of the disrupted gene.** Representative GMT islands (cyan rectangles) predicted to disrupt a gene upon insertion into the bacterial genome are shown above a predicted naive insertion site in a closely related *Vibrio* strain. The disrupted and the homologous genes are denoted with the same color. Gray rectangles denote homologous regions. The amino acid identity percentage between the protein encoded by the gene in which the predicted naïve insertion site is found and the homolog within the GMT island are denoted in blue rectangles. The strain names and the RefSeq accession numbers are provided.

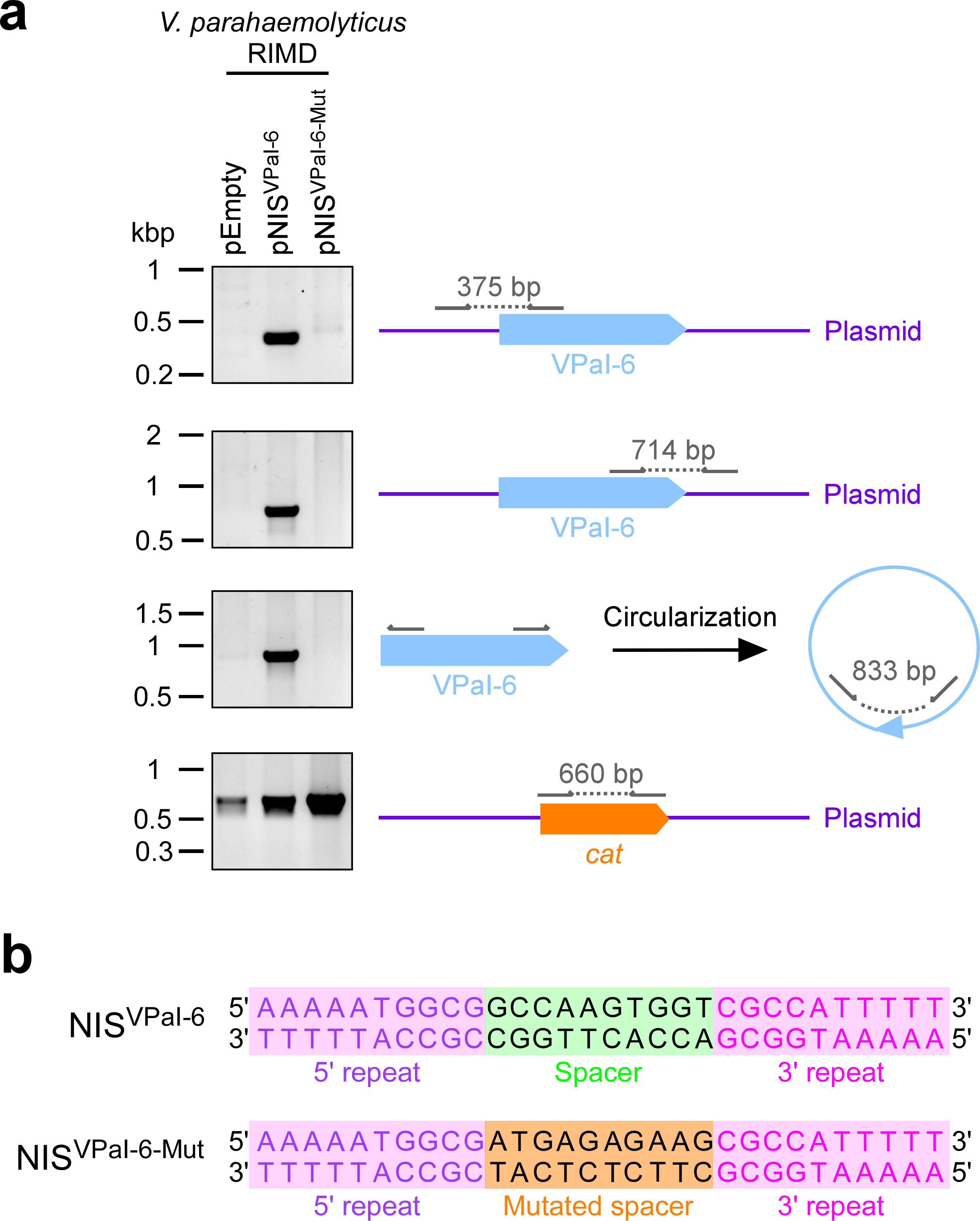

**Extended Data Fig. 3. The insertion site spacer sequence is important to facilitate GMT island insertion.** **(a)** Agarose gel electrophoresis analysis of the indicated amplicons. The total DNA isolated from wild-type *V. parahaemolyticus* RIMD 2210633 cells conjugated with an empty plasmid (pEmpty), a plasmid containing a predicted 30 bp-long naïve insertion site for VPaI-6 (pNIS^VPaI-6^), or a plasmid containing a mutated version of the insertion site in which the spacer between the inverted repeats was modified (pNIS^VPaI-6-Mut^), was used as a template. Arrows denote the positions of primers used for each amplicon; the expected amplicon size is denoted in gray. *cat*, chloramphenicol resistance gene found on the plasmids. **(b)** The sequences of the natural VPaI-6 naïve insertion site (NIS^VPaI-6^) and its mutated form (NIS^VPaI-6-Mut^) used in (a).

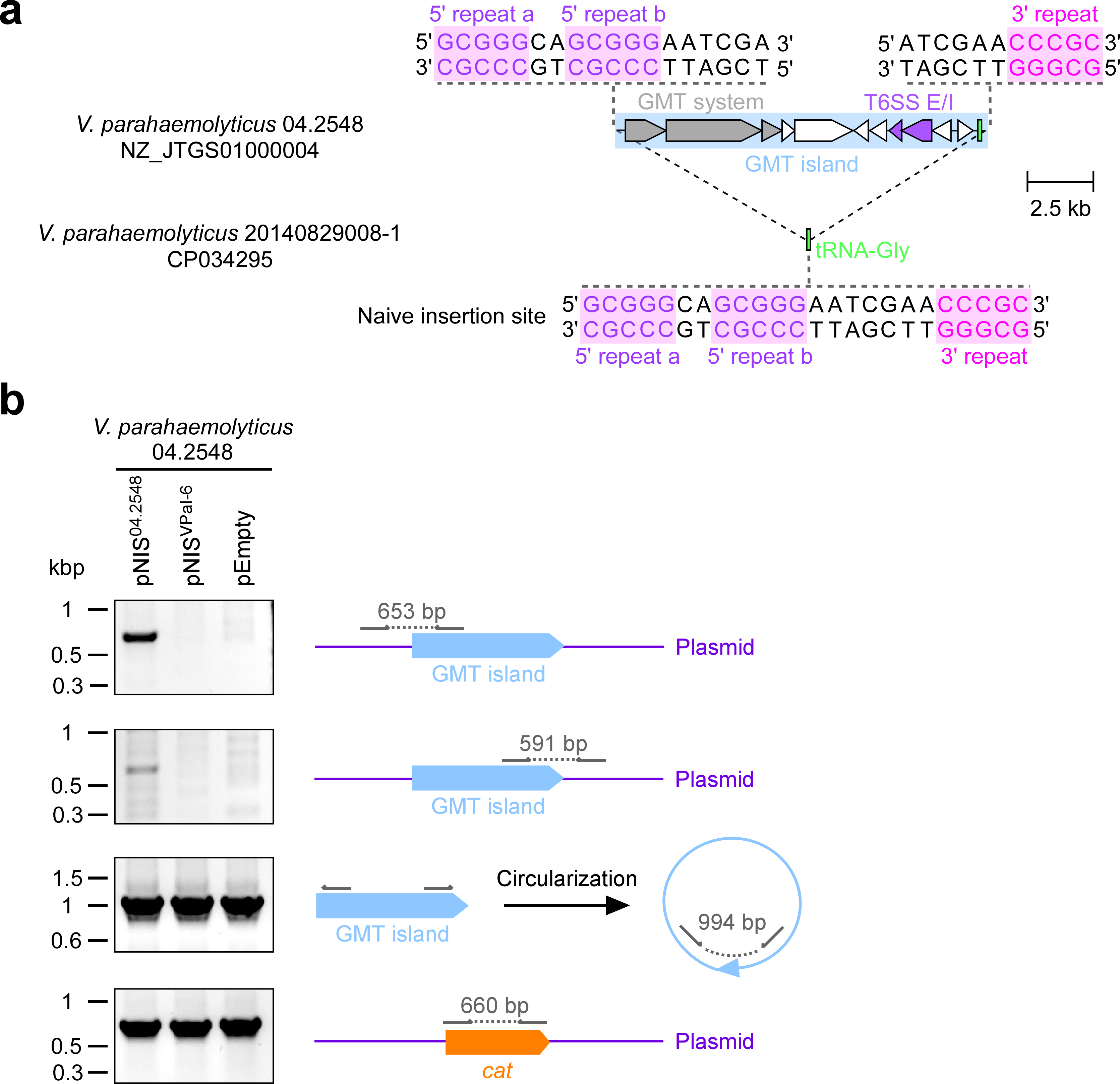

**Extended Data Fig. 4. The GMT island in *V. parahaemolyticus* 04.2548 is a functional mobile element.** **(a)** Schematic representation of the GMT island in *V. parahaemolyticus* 04.2548 (cyan rectangle). A predicted intragenic (tRNA-Gly), inverted repeat-containing (pink and purple-colored sequences) naïve insertion site identified in *V. parahaemolyticus* 20140829008-1 is shown below. Inverted repeat sequences flanking the GMT island are denoted. GenBank accession numbers are provided. **(b)** Agarose gel electrophoresis analysis of the indicated amplicons. The total DNA isolated from wild-type *V. parahaemolyticus* 04.2548 cells conjugated with an empty plasmid (pEmpty), a plasmid containing a predicted naïve insertion site for VPaI-6 (pNIS^VPaI-6^), or a plasmid containing a predicted naïve insertion site for the 04.2548 GMT island (pNIS^04.2548^), was used as a template. Arrows denote the positions of primers used for each amplicon; the expected amplicon size is denoted in gray. *cat*, chloramphenicol resistance gene found on the plasmids.

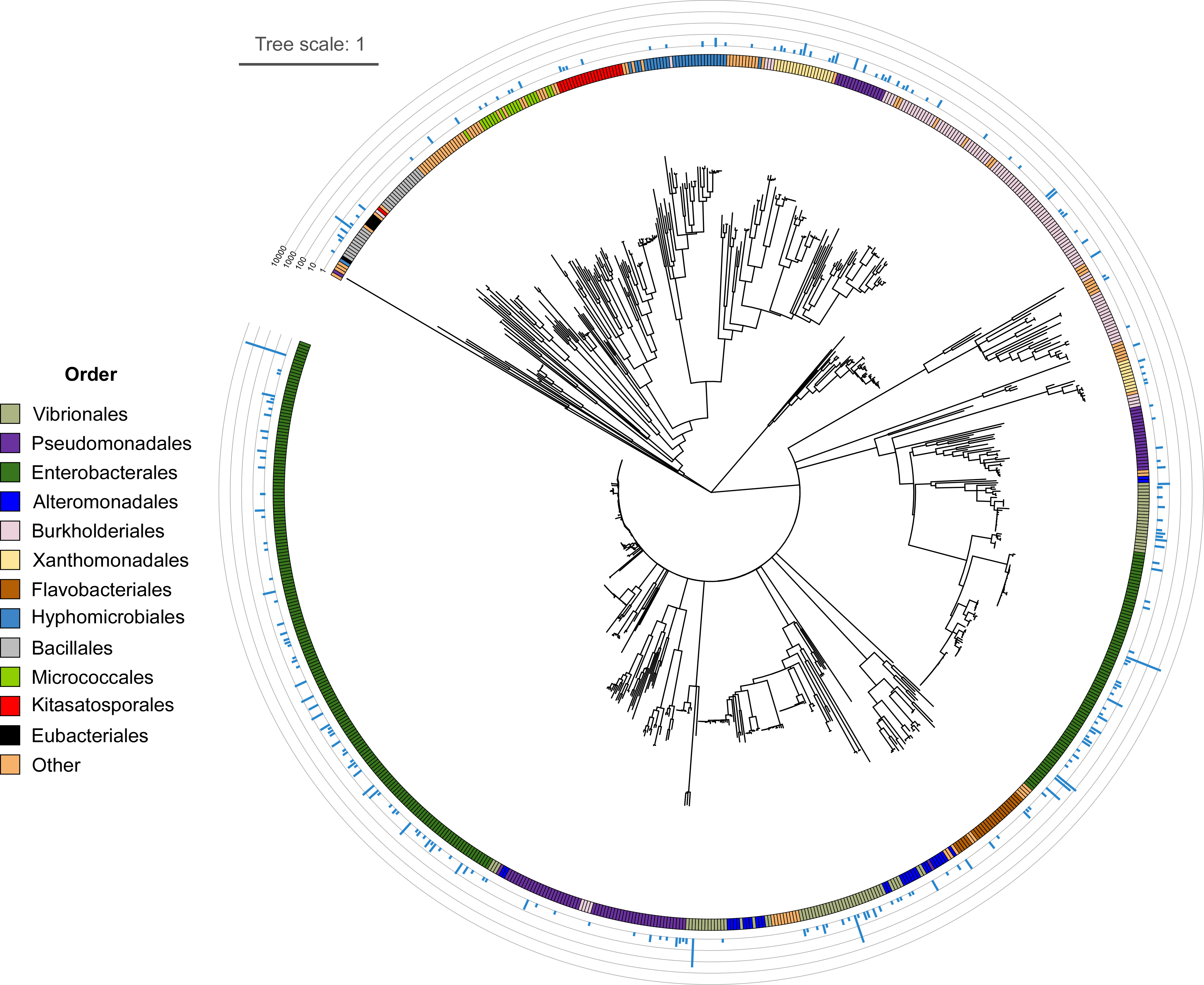

**Extended Data Fig. 5. GAPS1 homologs are widespread in bacteria.** Phylogenetic distribution of GAPS1 homologs. The bacterial order is denoted by color. Blue bars denote the number of genomes in which each protein accession was identified (Log_10_ scale). The evolutionary history was inferred by using the Maximum Likelihood method and LG+G+I+F model. The tree is drawn to scale, with branch lengths measured in the number of substitutions per site. Evolutionary analyses were conducted in MEGA X.

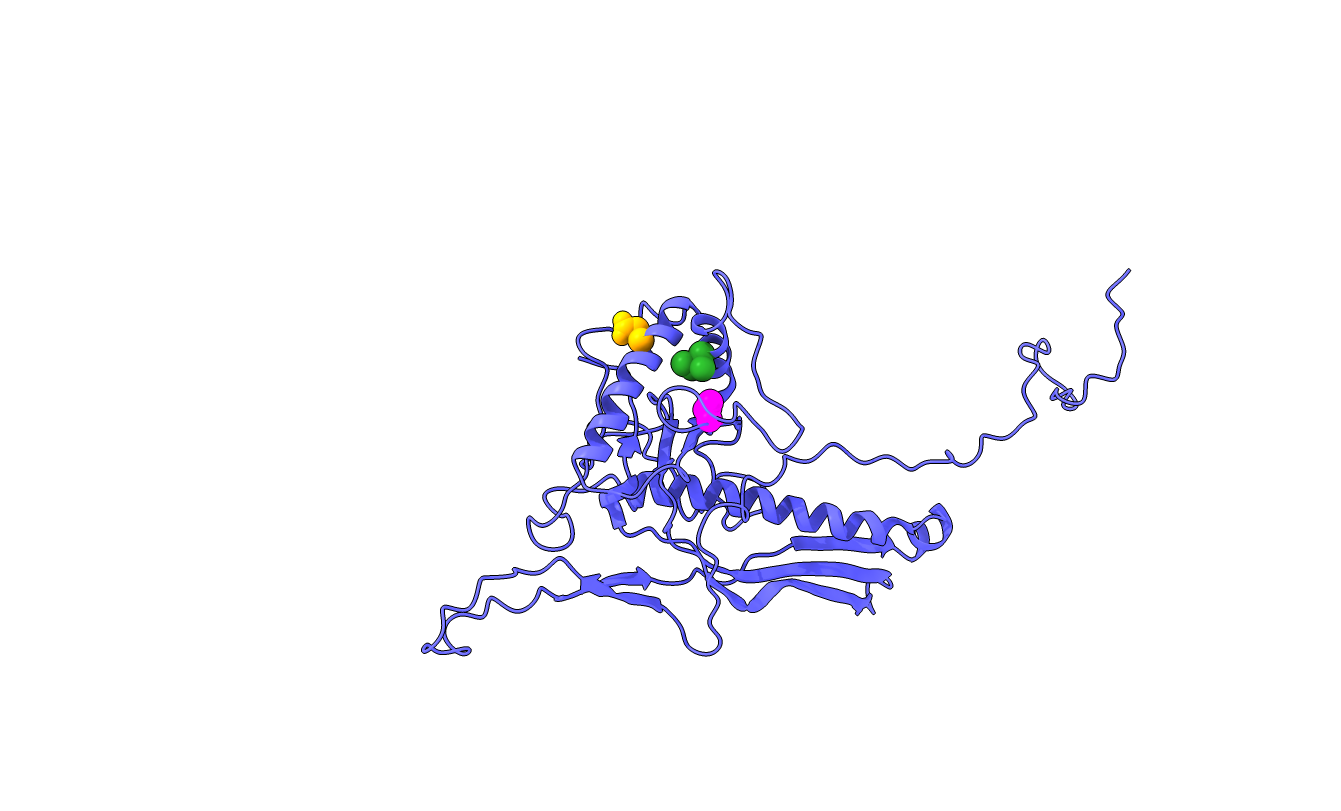

**Extended Data Fig. 6. Closely positioned residues in the T7 phage capsid protein are mutated in GAPS1 escape mutants.** The position of glutamic acid 183 (yellow), isoleucine 217 (green), and valine 247 (magenta) in the solved structure of the T7 phage capsid protein Gp10 (PDB:3j7x chain G).

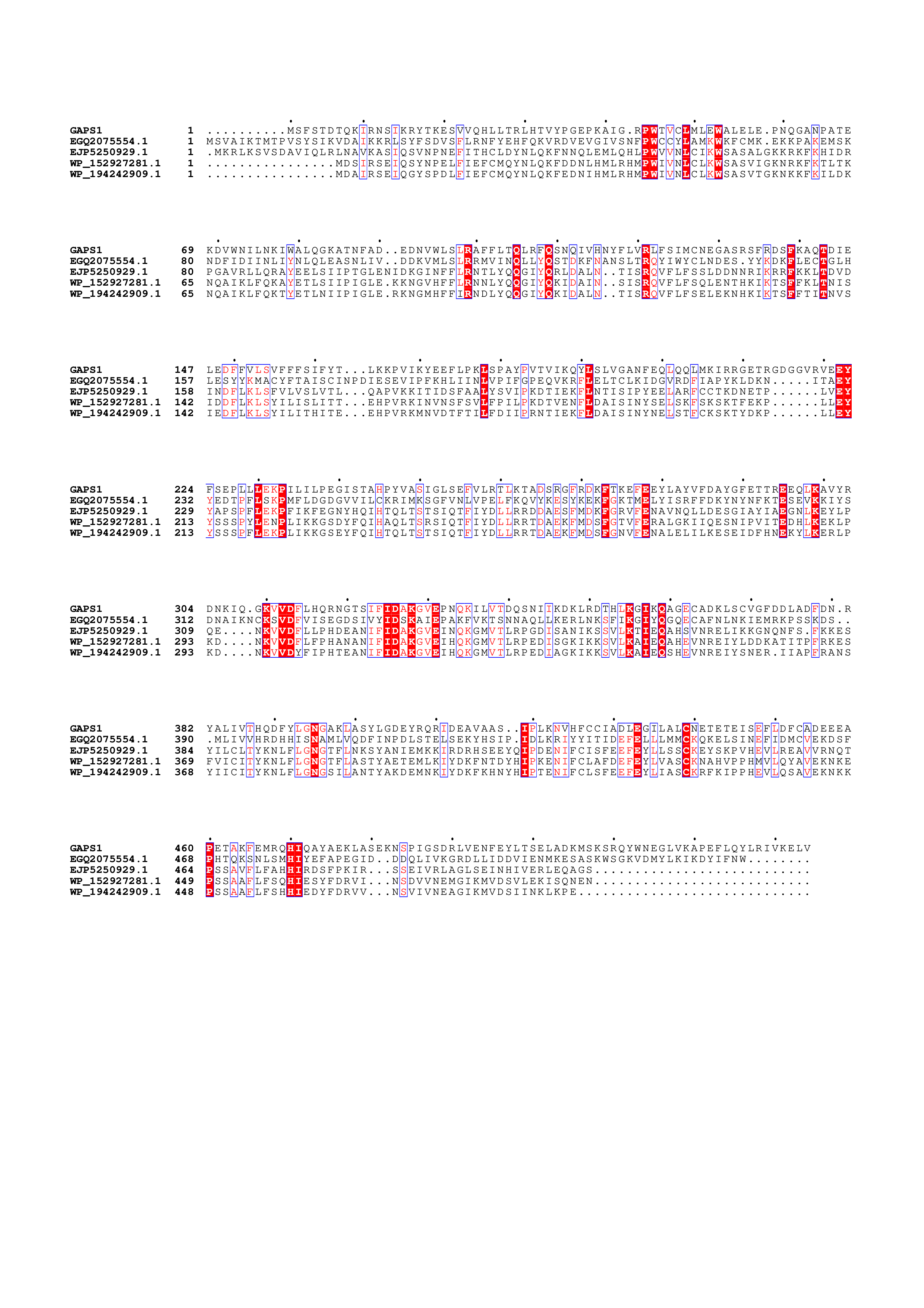

**Extended Data Fig. 7. GAPS1 homologs in *E. coli*.** Multiple sequence alignment for the indicated protein sequences was performed with Clustal W using Mega X. Similarity (red letter) and identity (red background) shading were done in ESPript 3.0^1^. The accession number of GAPS1 is WP_005477165.1.

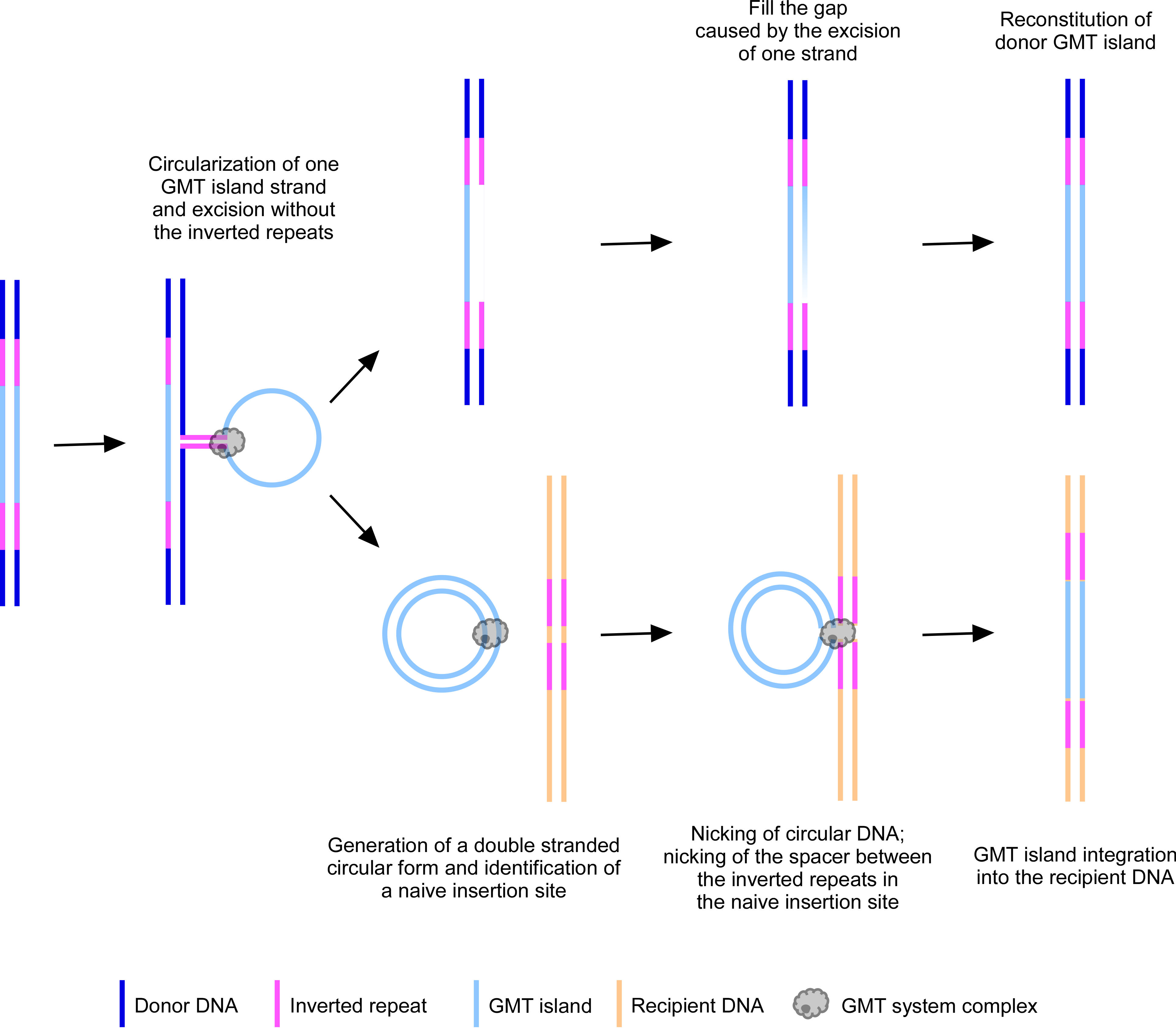

**Extended Data Fig. 8. A proposed model for GMT island replicative transfer.** A yet-unknown event or process leads to the excision and circularization of one of the GMT island strands without the flanking inverted repeat sequences; base pairing between the inverted repeat sequences may play a role in the process. A complementary strand is polymerized, resulting in a double-stranded circular form of the GMT island. The GMT system proteins, which are required for the excision and circularization, possibly remain bound to the junction between the ends of the GMT island and mediate recognition of a naïve insertion site containing a specific inverted repeat sequence. The naive insertion site is cleaved within the spacer sequence found between the inverted repeats, and the GMT island is inserted into the recipient site. In parallel, the cell replaces the excised strand in the donor DNA using the remaining strand as a template. This process results in two identical GMT islands.

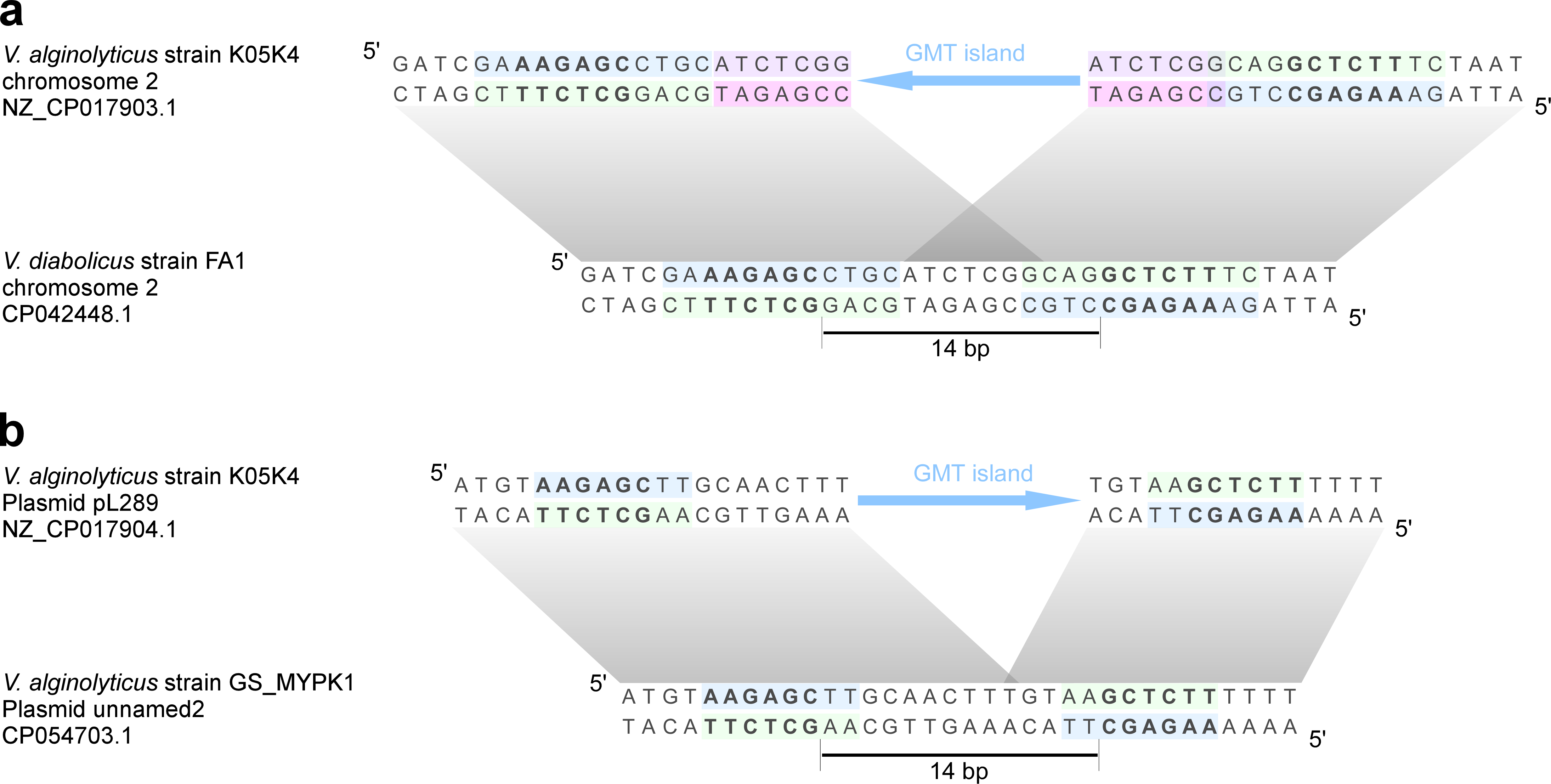

**Extended Data Fig. 9. Identical GMT islands on the chromosome and on a plasmid have similar predicted insertion sites. (a-b)** The nucleotide sequences flanking identical GMT islands on the chromosome (a) and on a plasmid (b) in *Vibrio alginolyticus*, and their predicted naïve insertion sites (shown below). Inverted repeat sequences are denoted in blue and green; the conserved repeat core found in both predicted insertion sites is denoted in bold. Direct repeat sequences resulting from apparent spacer duplication upon GMT island insertion into the chromosomal site are denoted in pink and purple (a). Gray rectangles denote identical sequences. The strain names and the GenBank accession numbers are provided.

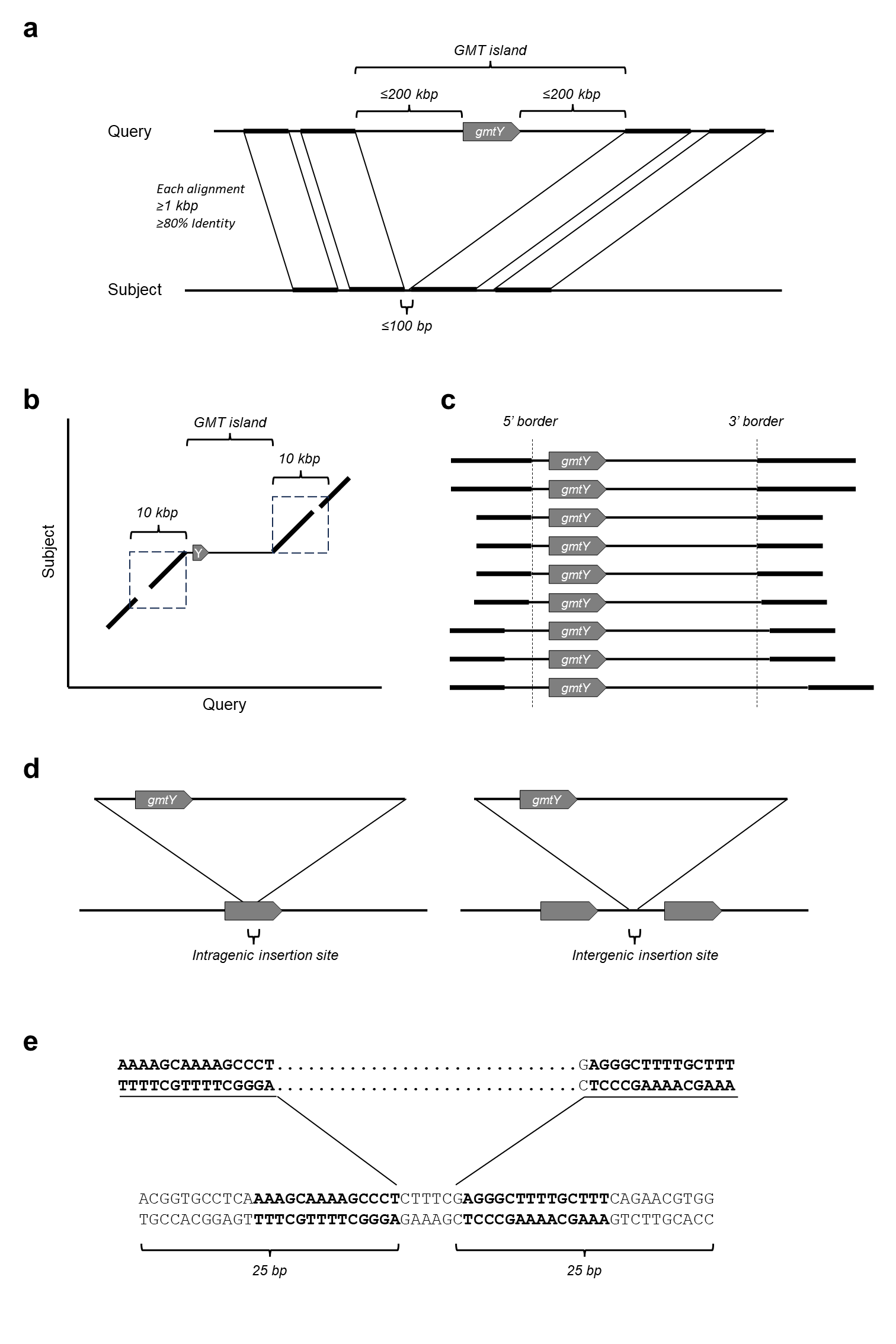

**Extended Data Fig. 10. Identification of GMT islands. (a)** Criteria for the identification of genomic accessions in closely related genomes that are homologous to the sequences flanking the GMT system. Thick lines represent the alignments between query and subject accessions. **(b)** Sequences upstream and downstream of the GMT islands should contain sequence alignments to the subject accessions in at least 4 kbp out of 10 kbp upstream and downstream sequences. **(c)** Grouping of alignments meeting all the requirements to determine the 5’ and 3’ borders of GMT islands. **(d)** Intragenic and intergenic insertion sites. **(e)** Analysis of sequences surrounding a predicted insertion site to identify direct and inverted repeats. Bold letters denote an inverted repeat found within the sequence example.

**Supplementary Tables (captions)**

**Table S1. Mutations found in GAPS1 T7 phage escape mutants.**

**Table S2. A list of bacterial strains used in this study.**

**Table S3. A list of gene fragments that were commercially synthesized for this study.**

**Table S4. A list of plasmids used in this study.**

**Table S5. A list of primers used in this study.**

**Table S6. A list of bacteriophages used in this study.**

**Supplementary Datasets (captions)**

**Dataset S1. A list of GMT proteins and adjacently encoded proteins.**

**Dataset S2. The borders of GMT islands found within complete genomes, and the presence of a T6SS in these strains.**

**Dataset S3. Anti-phage defense systems and antibacterial T6SS effectors identified in GMT island cargoes.**

**Dataset S4. A list of GAPS1 homologs.**

**Supplementary Files (captions)**

**File S1. Sequencing results of PCR-amplified DNA for data shown in Fig. 1d.**

**File S2. Sequencing results of PCR-amplified DNA for data shown in Extended Data Fig. 4b.**
