## Supplementary material for "Gamma-Mobile-Trio systems define a new class of mobile elements rich in bacterial defensive and offensive tools": File S1

| **Mobility product** | **Sanger sequencing (5'-3')** | **Primer used for sequencing** |
| --- | --- | --- |
| VPaI-6  5'-junction with plasmid | 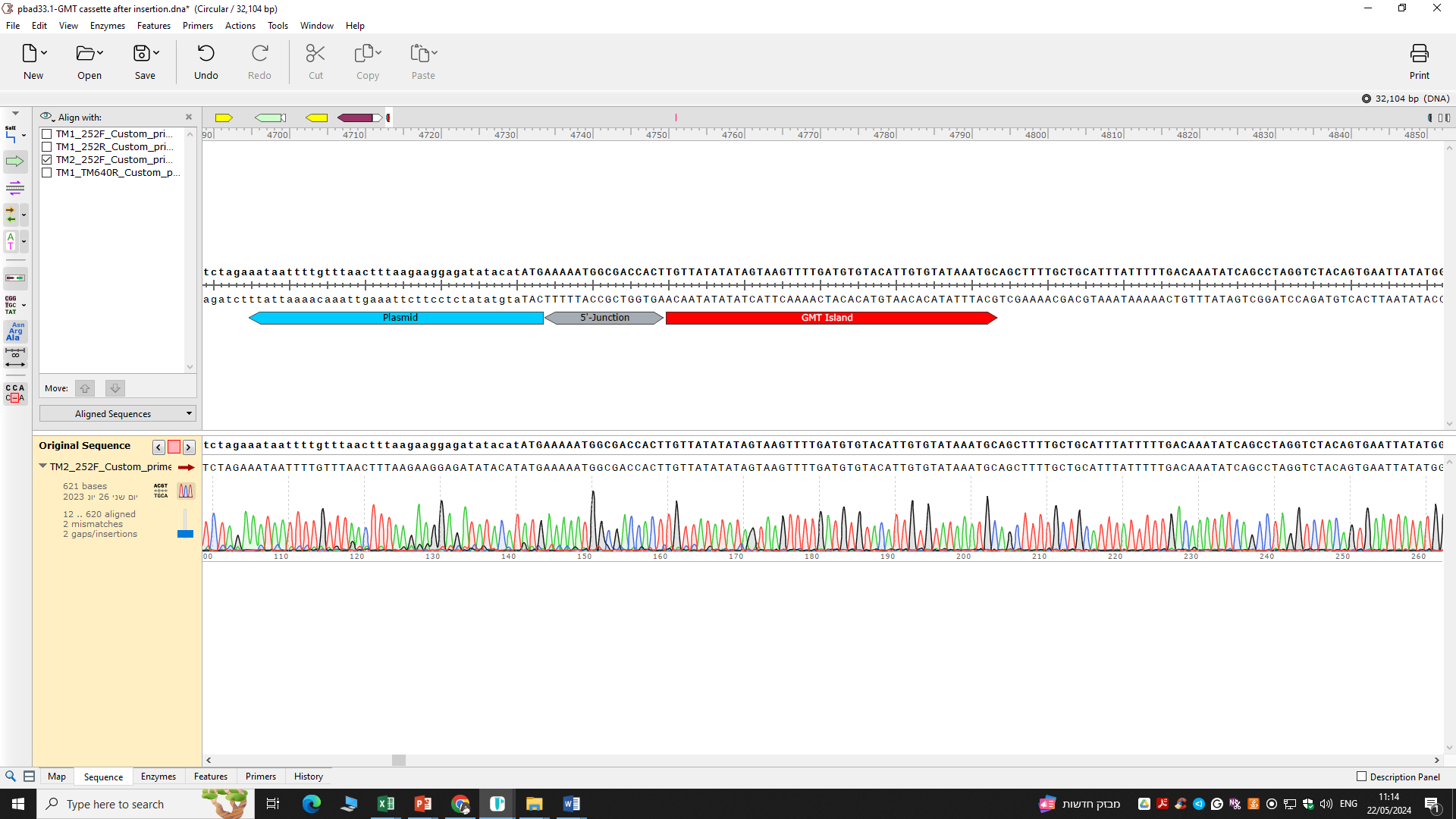 | 252F |
| VPaI-6  3'-junction with plasmid | 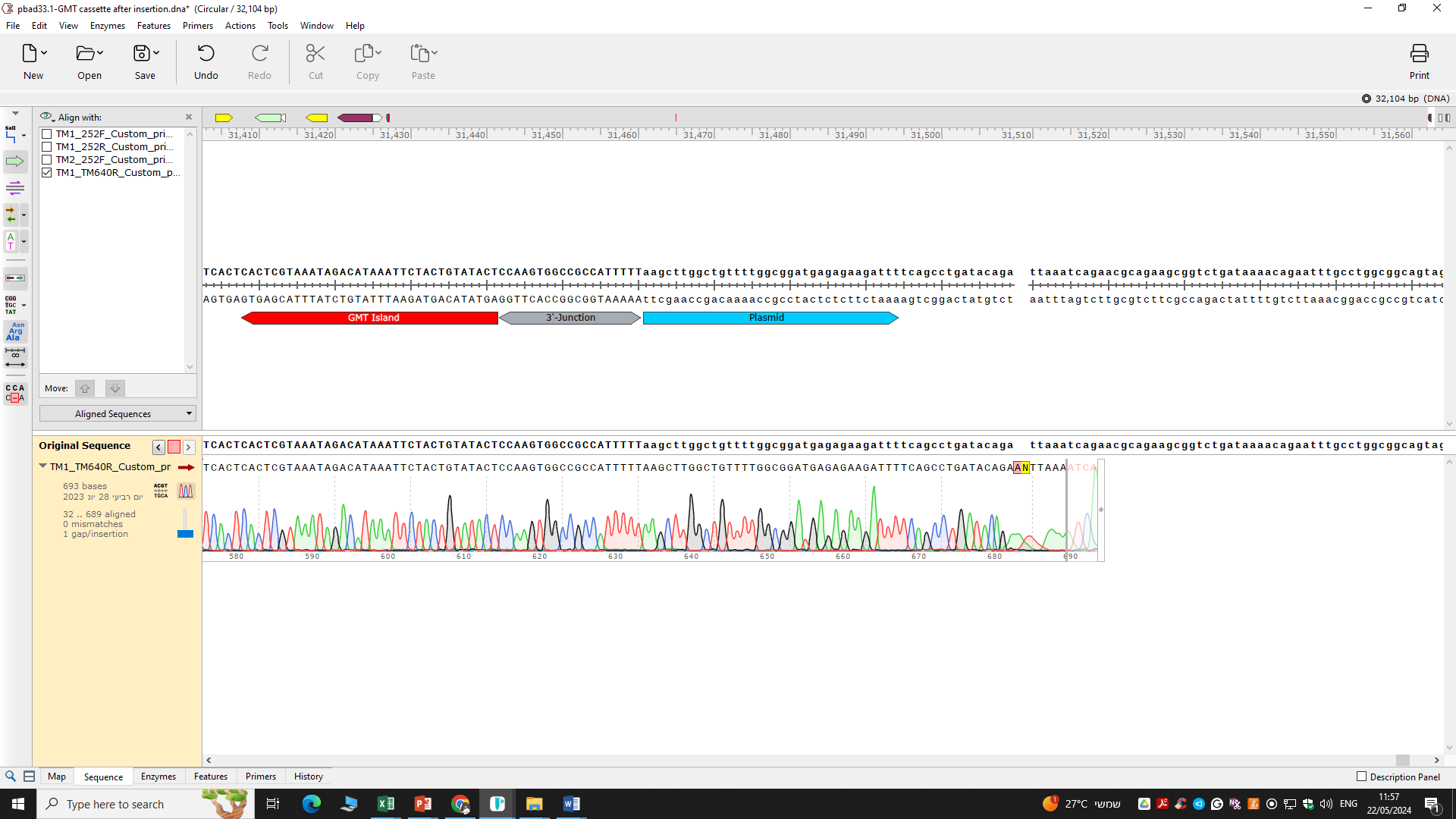 | TM640R |
| Circular VPaI-6 | 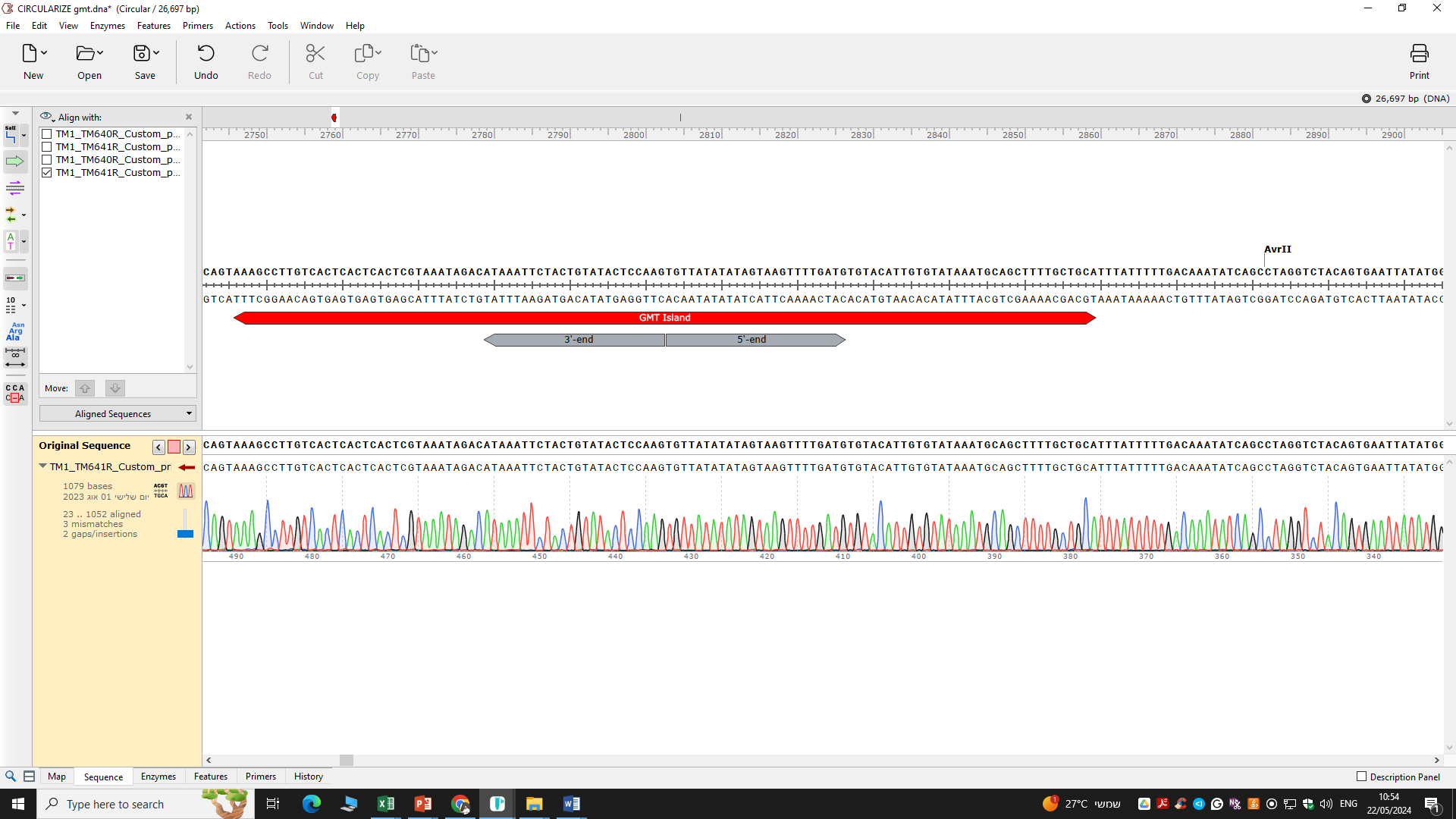 | TM640R |
