## Supplementary material for "Gamma-Mobile-Trio systems define a new class of mobile elements rich in bacterial defensive and offensive tools": File S2

| **Mobility product** | **Sanger sequencing (5'-3')** | **Primer used for sequencing** |
| --- | --- | --- |
| 04-2548  5'-junction with plasmid | 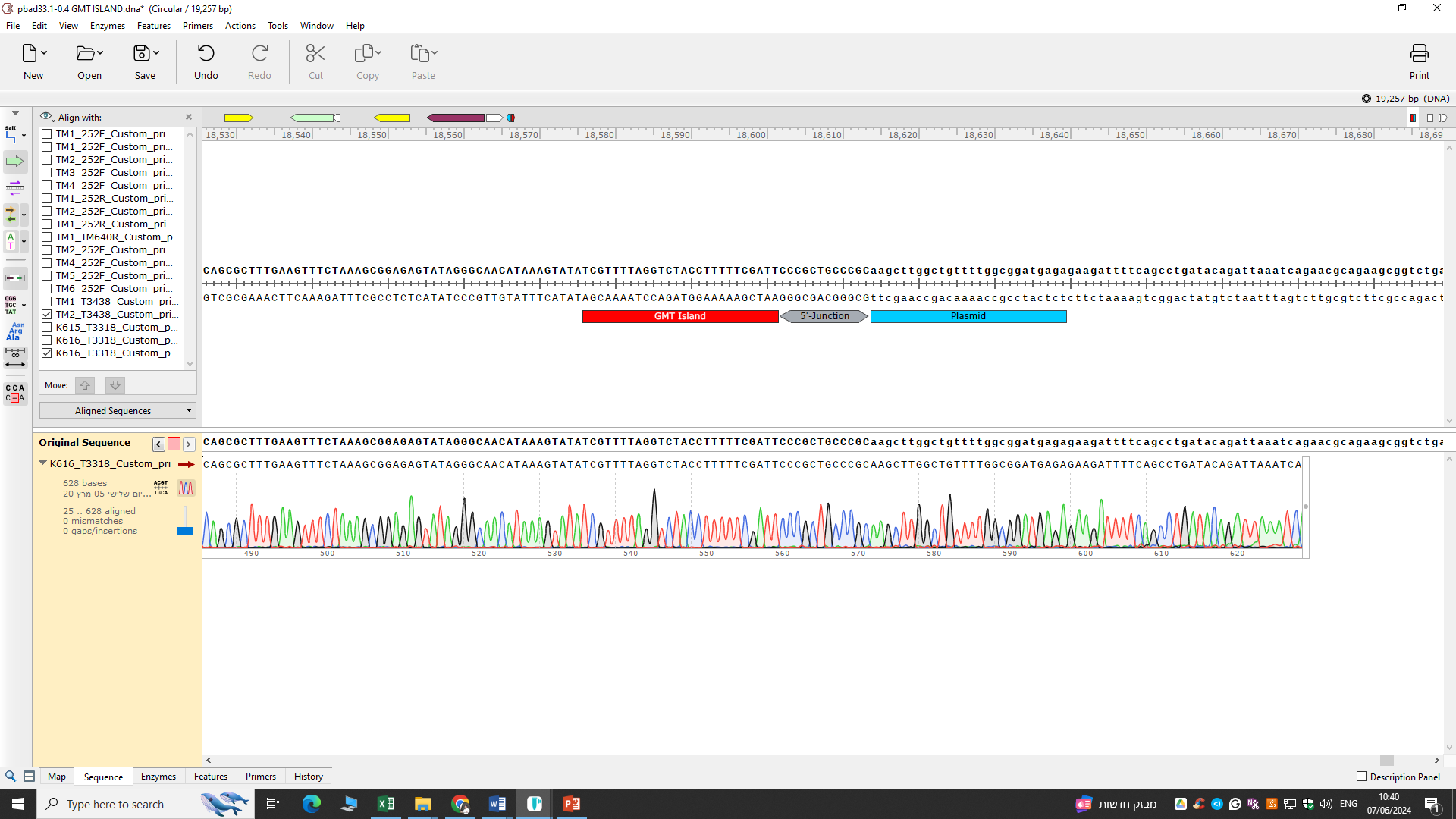 | T3318 |
| 04-2548  3'-junction with plasmid | 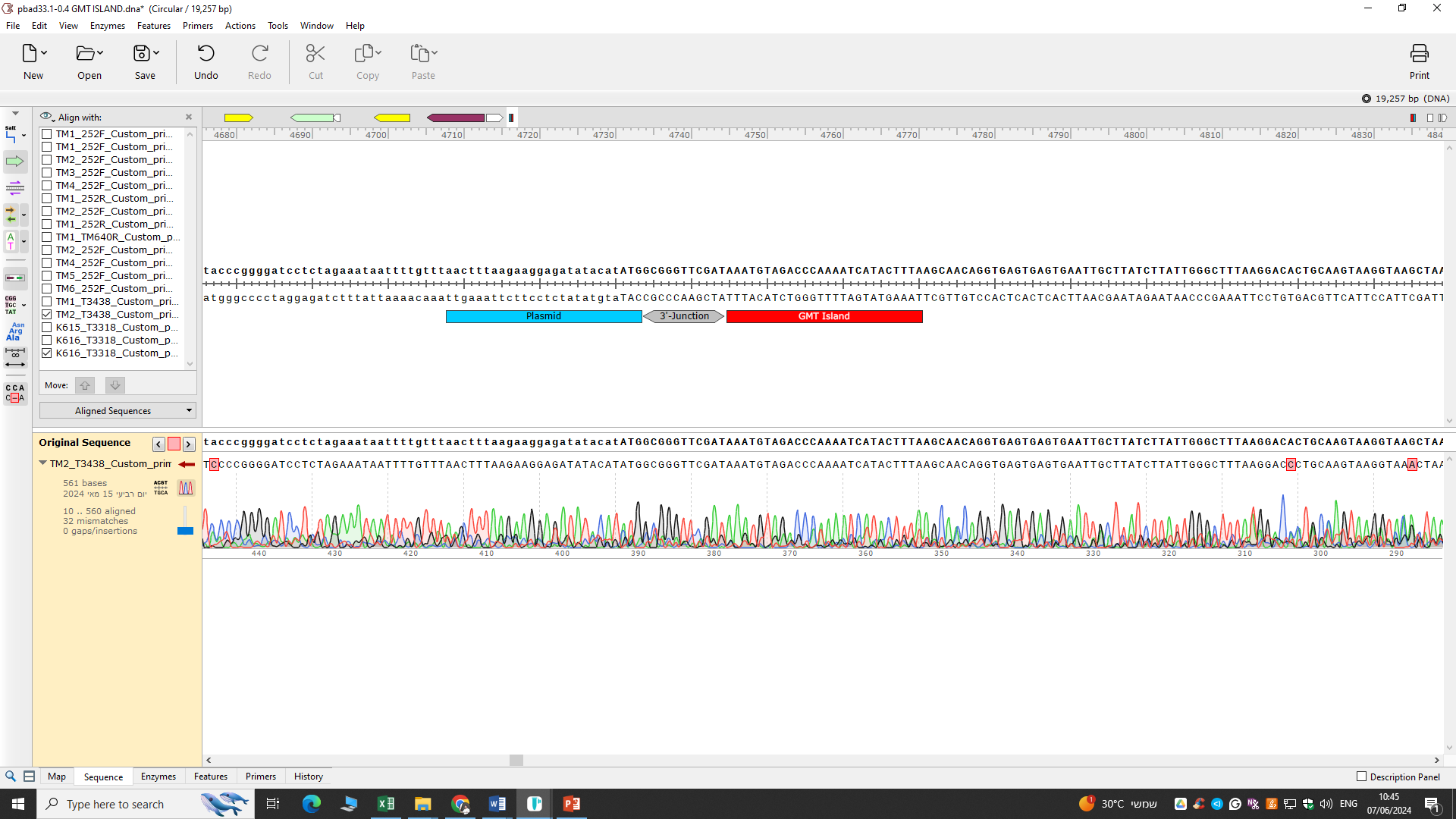 | T3438 |
| Circular  04-2548 | 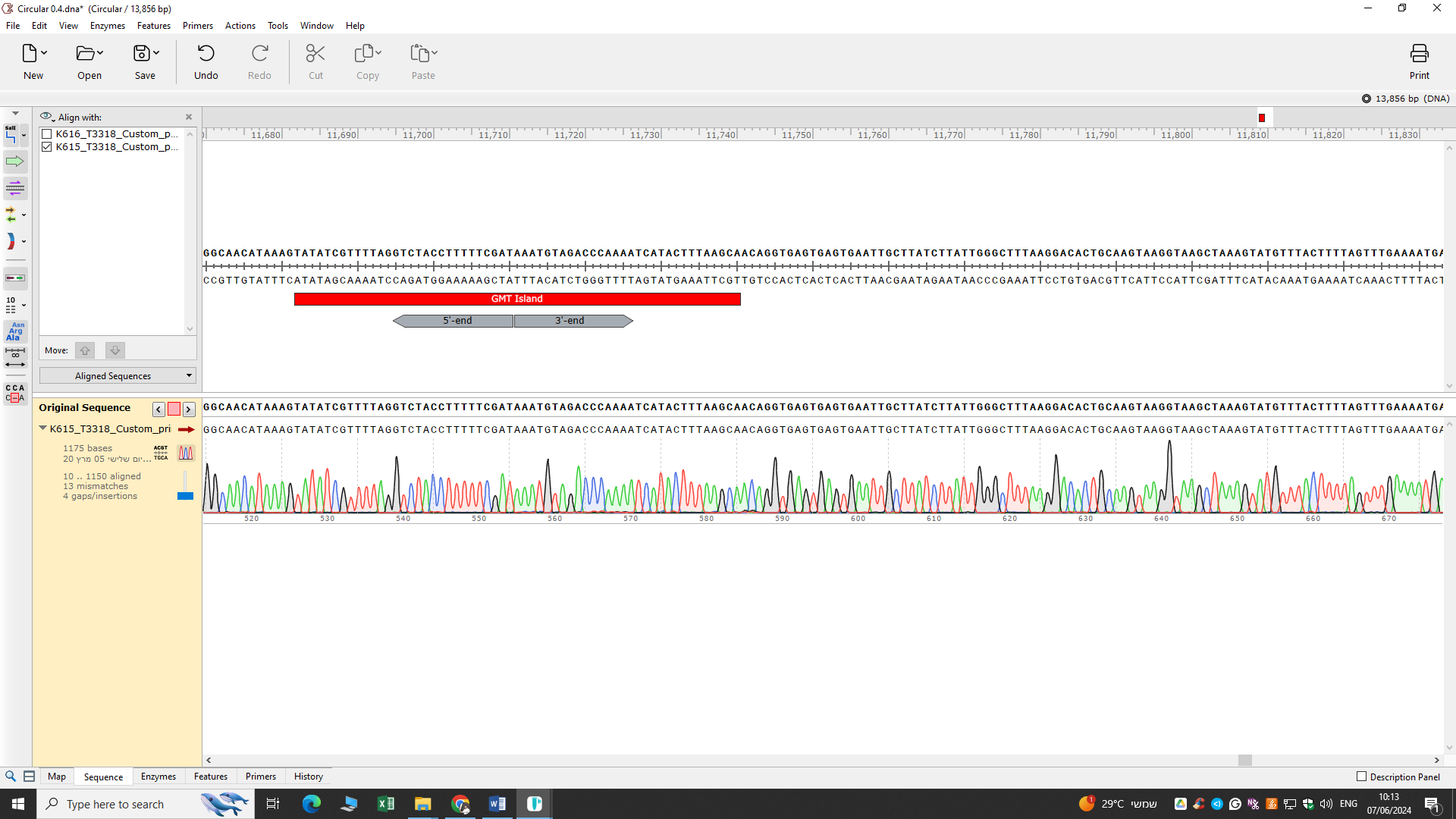 | T3318 |
